# Endothelial-fibroblast interactions during Scarb1 accelerate heart failure

**DOI:** 10.1101/2023.09.15.557661

**Authors:** Toshiomi Katsuki, Dai Kusumoto, Yohei Akiba, Mai Kimura, Jin Komuro, Takahiro Nakamura, Hisayuki Hashimoto, Thukaa Kuoka, Yutaka Suzuki, Yoshiaki Kubota, Keiichi Fukuda, Shinsuke Yuasa, Masaki Ieda

**Author notes:** Corresponding author: Dai Kusumoto, M.D., Ph.D., Center for Preventive Medicine / Department of Cardiology, Keio University School of Medicine, 35 Shinanomachi Shinjuku-ku, Tokyo 160-8582, Japan, or Shinsuke Yuasa, M.D., Ph.D., Department of Cardiology, Keio University School of Medicine, 35 Shinanomachi Shinjuku-ku, Tokyo 160-8582, Japan.

## Abstract

Endothelial cells (ECs) maintain cardiac homeostasis and EC dysfunction causes heart failure progression. Moreover, pathological changes occur via interactions between multiple cells, including ECs. Here, we conducted single-cell RNA-seq analysis of non-cardiomyocytes in mouse hearts during heart failure progression to elucidate the pathological changes in ECs and fibroblasts (FBs) mediated by cell-cell interactions. We show that capillary and arterial ECs exhibit mesenchymal gene expression changes with heart failure progression, indicating that endothelial-to-mesenchymal transition (EndMT) is a major pathological alteration in ECs. We also found that the interaction between ECs and FBs was enriched during heart failure, particularly when involving Scavenger Receptor Class B Member 1 (Scarb1) in ECs. FBs induce mesenchymal gene alterations in ECs in the EC-FB co-culture system, which is inhibited by blocking SCARB1. RNA-seq analysis showed that administration of a SCARB1 inhibitor blocked mesenchymal gene expression, and inflammatory changes, suggesting that the EC-FB interaction via Scarb1 is important for EndMT induction in ECs. Systemic administration of a SCARB1 inhibitor attenuated heart failure progression and cardiac fibrosis. EC-specific *Scarb1* knockout mouse showed improved cardiac function, suggesting a crucial role of Scarb1 in heart failure progression. Our results suggest that Scarb1 is a promising candidate for novel heart failure treatments that target ECs.

## Introduction

Heart failure significantly contributes to shortened global healthy life expectancy ^1^. Although there are various cell types in the heart, previous studies have focused on cardiomyocytes, given their crucial roles including pumping blood in the heart^2–5^. Non-cardiomyocytes (non-CM), which comprise nearly 70% of the cells in the heart^6^, have been shown to significantly affect the progression of heart failure^7–9^. Among them, the endothelial cell (EC) population is the largest^6^. ECs play important roles in maintaining cardiac homeostasis^10–12^, and EC dysfunction has been contributes to the progression of heart failure^13–15^. As EC-specific gene modifications improved heart failure progression^16–18^, they are a potent therapeutic target for heart failure. However, the detailed mechanisms of EC dysfunction and its contribution to heart failure progression remain unclear. Furthermore, in the progression of heart failure, pathological changes in cells occur at the individual cells level and through interactions between multiple cells, including ECs^19–22^. Therefore, instead of solely analysing individual cell populations, investigating how multicellular interactions change during the progression of heart failure is key. Single-cell RNA sequencing (scRNA-seq) enables a comprehensive analysis of the transcriptional profiles of various cells within tissues at the single-cell level and can reveal cell-to-cell interactions by calculating ligand-receptor pair expression^23–25^.

In this study, we performed scRNA-seq analysis targeting non-CMs through heart failure progression to elucidate the pathological changes in cells, mainly focusing on ECs and FBs and their pathological cell-cell interactions. Furthermore, we focused on the Scavenger Receptor Class B Member 1 (Scarb1) expressed in ECs based on cell-to-cell interaction analysis and proposed that Scarb1 could be a novel therapeutic target for heart failure.

## Results

### Overview of non-CMs in the heart

To investigate the pathological changes in non-CMs during heart failure development, we created transverse aortic constriction (TAC) mouse models^26,27^. We extracted non-CM from hearts in the normal phase (TAC 0 week), adaptive hypertrophy phase (TAC 2 weeks), and heart failure phase (TAC 12 weeks) in TAC model mice (Extended Data Fig. 1a,b) and conducted droplet-based single-cell RNA-sequence analysis (scRNA-seq)^28^ (Fig. 1a,b, and Extended data Fig. 1c,d). We integrated time-series data from the three phases of heart failure development using gene anchors^29^ and identified cell types based on known gene expression (Fig. 1c, d). The machine learning-based classification algorithm successfully classified the same cell types in different heart failure stages into the same clusters (Fig. 1f). Endothelial cells (ECs), fibroblasts (FBs), blood cells (BCs), smooth muscle cells (SMCs), and neural cells (NCs) were identified in descending order of cell number, and CMs were not included (Extended data Fig. 1e, f). The number of ECs increased with the progression of heart failure, especially in the hypertrophic phase (Fig. 1e), indicating that the disruption of homeostasis by ECs plays a pivotal role in heart failure progression. We then re-clustered the major populations into subpopulations to analyse the detailed pathological changes in each cellular type during heart failure development.

**Fig. 1:**
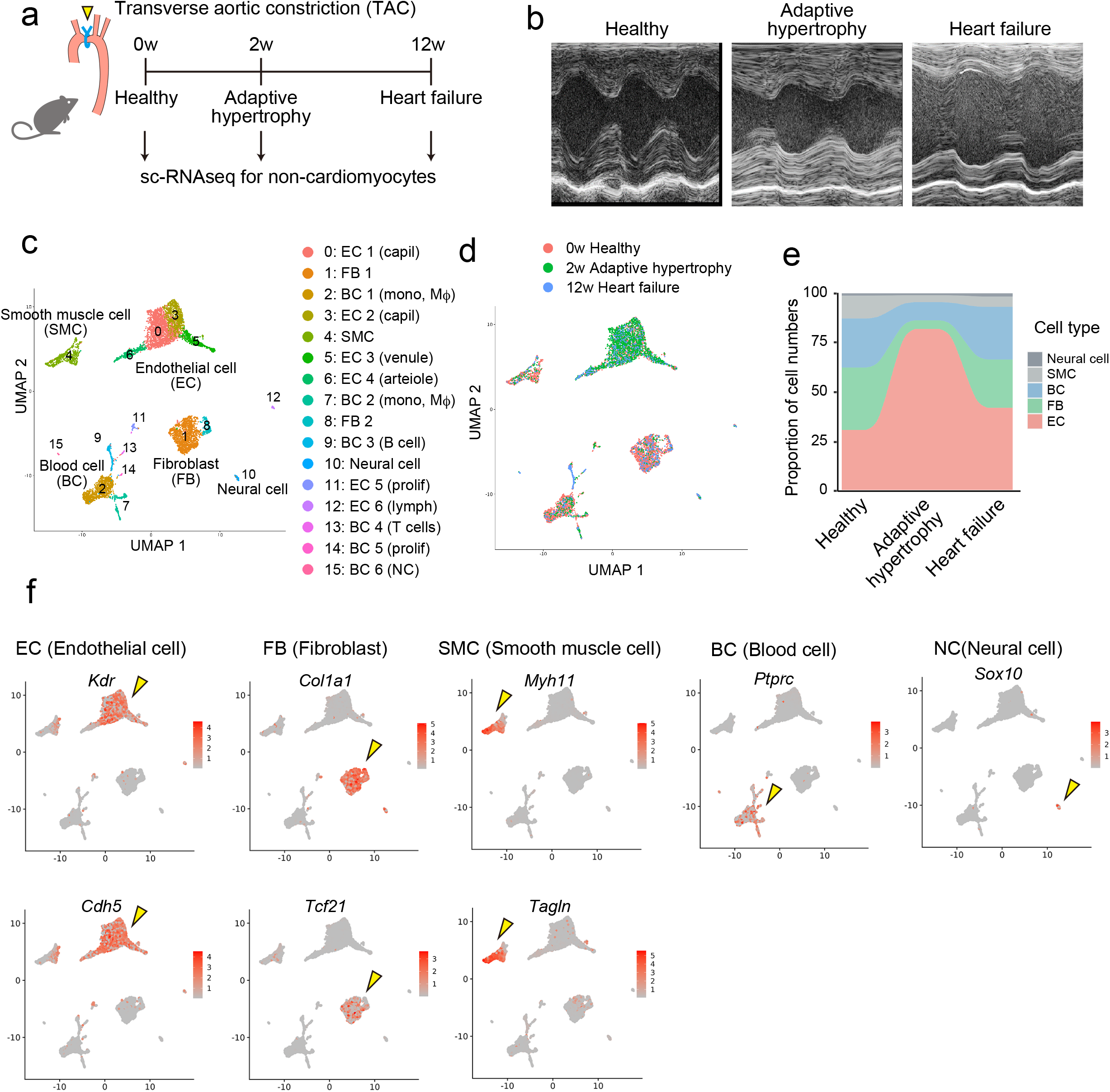
Overview of non-cardiomyocytes via heart failure progression analysed by Single-cell RNA-sequence (scRNA-seq). a, Experimental protocol for scRNA-seq analysis. Healthy, adaptively hypertrophic, and failed murine hearts were subjected to scRNA sequencing. Two weeks and twelve weeks after the TAC procedure, adaptive hypertrophy and heart failure phases occurred, respectively. **b,** Representative M-mode echocardiograms of sacrificed mice. The healthy (left), adaptive hypertrophy (middle), and heart failure (right) phases are displayed. **c,** Visualization of single-cell transcriptomes from combined datasets of healthy, adaptive hypertrophic, and failed murine heart samples in a UMAP dimensional reduction manner (UMAP plot). Each point corresponds to a single-cell dataset. The subgroups are shown in different colours. **d,** Visualization of the same data in the previous plot (c) according to the time course. **e,** Proportional changes in crude cell types over time The x-axis represents time, and the y-axis represents the proportions of EC, FB, BC, SMC, and neural cells in the total number of cells. **f,** Visualization of cell type-specific UMI counts for EC, FB, SMC, BC, and neurons. Richer red indicated more abundant expression. Yellow arrowheads indicate the most specific subgroup. TAC, transverse aortic constriction; UMAP, uniform manifold approximation and projection; EC, endothelial cells; FB, fibroblasts; BC, blood cells; SMC, smooth muscle cells.

### Pathological changes in fibroblasts through heart failure progression

Among the five major clusters, we first focused on FB subpopulations because validation of FB pathology may confirm the validity of our scRNA-seq analysis. By reclustering the FB population, they were divided into seven sub-clusters (Fig. 2a). Pseudo-time analysis ^30^ of the FB clusters showed that cluster 0 was the starting point of the pseudo-time trajectories, and the stem trajectory diverged into cluster 5,1 direction and cluster 2,4 direction with the progression of heart failure (Fig. 2b). Therefore, cluster 0 was considered as healthy FBs, which was abundant in the healthy heart, namely, resident FBs. In contrast, Clusters 1 and 4 were considered pathological FBs, which were abundant in the diseased heart and located at the ends of the trajectories. Regarding cell numbers, Cluster 1 increased during adaptive hypertrophy, whereas Cluster 4 increased during the heart failure (Fig. 2c). Extracellular matrix (ECM) production including periostin (*Postn*) in FB increased with heart failure progression, and it have been identified as key features of pathological FB changes in heart failure ^31,32^. Our analysis revealed that clusters 1 and 4 were *Postn*-positive (Fig. 2d), suggesting that these FB clusters are potentially harmful to the progression of heart failure. Moreover, FBs in cluster 4 expressed stress fibres such as *Acta2* (Fig. 2e), a unique gene in myofibroblasts (myoFBs) ^33–35^. Thus, Cluster 1 was identified as an activated FB population that was *Postn*-positive and *Actb*-negative, whereas Cluster 4 was identified as a myoFB population that was *Postn*-positive and *Actb*-positive. Conversely, the expression of genes characteristic of cardiac FBs, such as *Tcf21* and *Pdgfra*, was decreased in myoFBs, suggesting that FBs were undergoing a transformation (Fig 2f). According to GO term analysis, genes related to the ECM and collagen were enriched in activated FB (Fig. 2g), whereas genes related to bone development, Tgf-α, and Wnt signalling were enriched in myoFBs (Fig. 2h). These findings suggest that both clusters are pathological FBs implicated in heart failure progression, and our single-cell analysis accurately captured their changes.

**Fig. 2:**
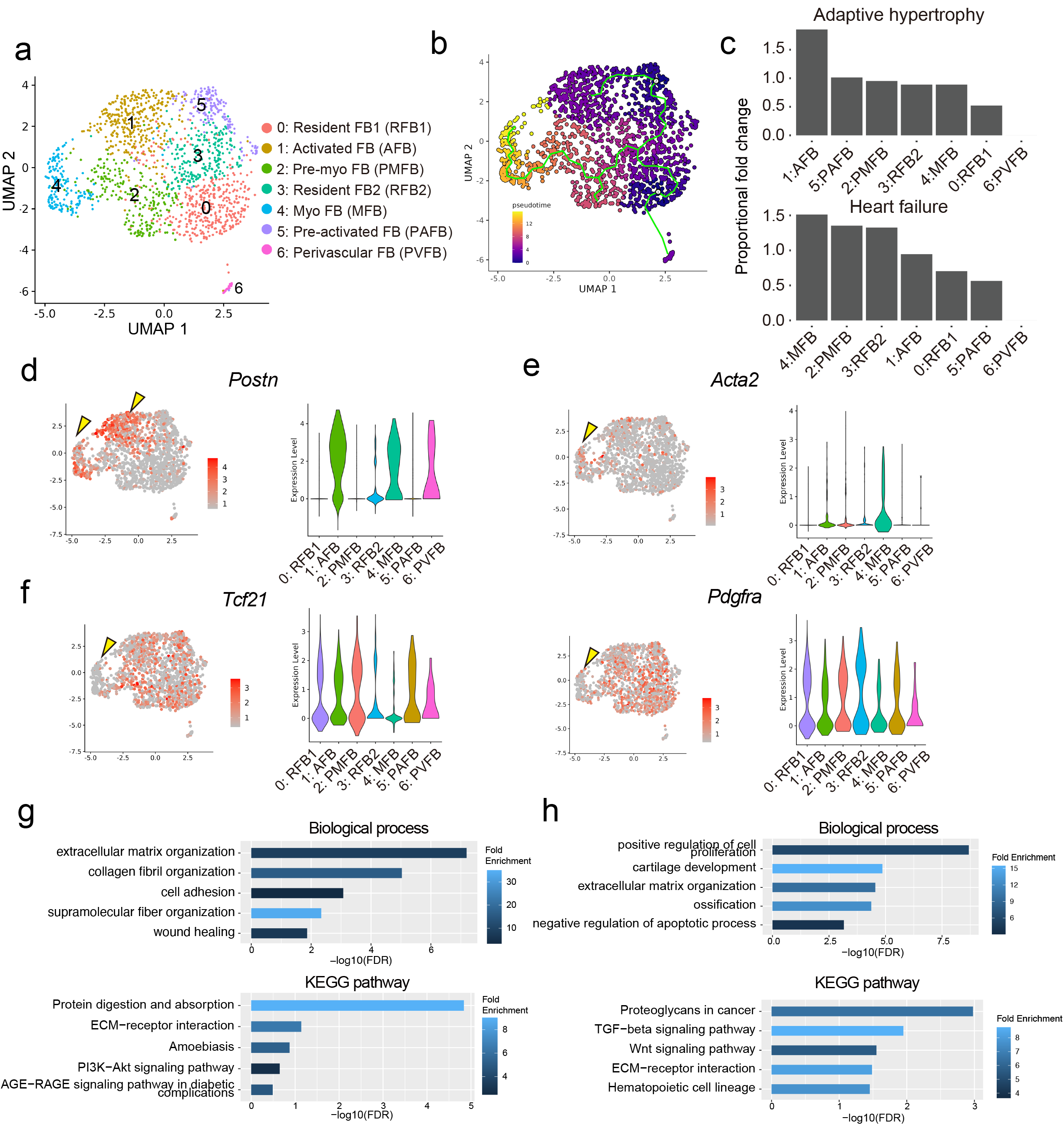
scRNA-seq captured gene expression profiles of classical and unknown fibroblasts subtypes through heart failure development. a, UMAP plot for fibroblasts in combined datasets of healthy, adaptively hypertrophic, and failed hearts. The cells were re-clustered before visualization. **b,** Pseudotime overlayed UMAP plot. Pseudotime was calculated along gene expression similarity of cells. Green lines represent the cell trajectories. Trajectories were aligned using similarity measures, indicating the virtual differentiation of the cells. Brighter colours indicate more differentiated cells. **c,** Proportional fold-changes of each subgroup cell number in adaptive hypertrophy and heart failure compared to the healthy state. The subgroups were arranged in descending order of fold-change. **d-f,** Visualization of key fibroblast gene expression markers for (d) activated fibroblasts, (e) myofibroblasts, and (f) common cardiac fibroblasts. Yellow arrowheads indicate the characteristic gene expression patterns of activated cells and myofibroblasts. The distributions of gene expression levels in the subgroups were displayed using violin plots. **g-h,** Visualization of GO analysis and KEGG pathway analysis. Rows are top-5 significant GO terms for biological processes/KEGG pathways in (g) activated fibroblasts and (h) myofibroblasts. X axis was −log10(FDR). A brighter colour indicates a higher fold enrichment. UMAP, uniform manifold approximation and projection; GO, gene ontology analysis; KEGG, Kyoto Encyclopedia of Genes and Genomes; FDR, false discovery rate.

### Pathological changes in endothelial cells through the heart failure progression

We then analysed the changes in ECs, the largest cell population in the heart, during the progression of heart failure. We reclustered the ECs and classified them into 11 cell populations (Fig. 3a). Using differential gene expression analysis, ECs were classified as capillary EC (CEC 1,2,3,4), fibrotic CEC, arterial EC (AEC), arteriole EC (ALEC), venous EC (VEC), venule ECs (VLEC), proliferative EC (PEC), and lymphatic endothelial cells (LEC) (Fig. 3b). By analysing the clusters that increased in number during heart failure progression, we found that cluster 6 increased during the period of adaptive hypertrophy, whereas cluster 5 increased during the heart failure phase. Cluster 6 was a type of capillary EC based on the expression of factors related to lipid metabolism^36,37^, such as *Cd36* and *Fabp4* (Fig. 3b). Pseudotime analysis revealed that cluster 6 differentiates from Cluster 0, a common capillary EC in healthy hearts. GO term analysis showed that the expression of genes related to the ECM, especially collagen, was elevated in cluster 6. Violin plot analysis also revealed increased expression of *Col1a1* and *Col3a1* (Fig. 3f). These genes are characteristic of mesenchymal cells; Thus, Cluster 6 cells can be defined as ECs undergoing endothelial-mesenchymal transition (EndMT)^38,39^ to acquire mesenchymal cell traits, namely fibrotic capillary EC. *Il1α* was also upregulated in fibrotic capillary EC (Fig. 3). Cluster 5, which increased in number during the heart failure phase, exhibited arterial EC characteristics. When clusters were re-analysed by heart failure developmental phase, we found that arterial EC could be divided into two groups: healthy and heart failure phases. GO analysis showed that gene expression related to the actin cytoskeleton was enriched. Stress fibre-related genes, such as *Acta2* and *Tagln* were upregulated in arterial ECs during the heart failure phase. *Fgfr1*, which plays an important role in organ fibrosis, is also upregulated in arterial ECs during heart failure, indicating its contribution to heart fibrosis. These findings suggest EndMT is a key characteristic change in both capillary ECs and arterial ECs during the heart failure progression.

**Fig. 3:**
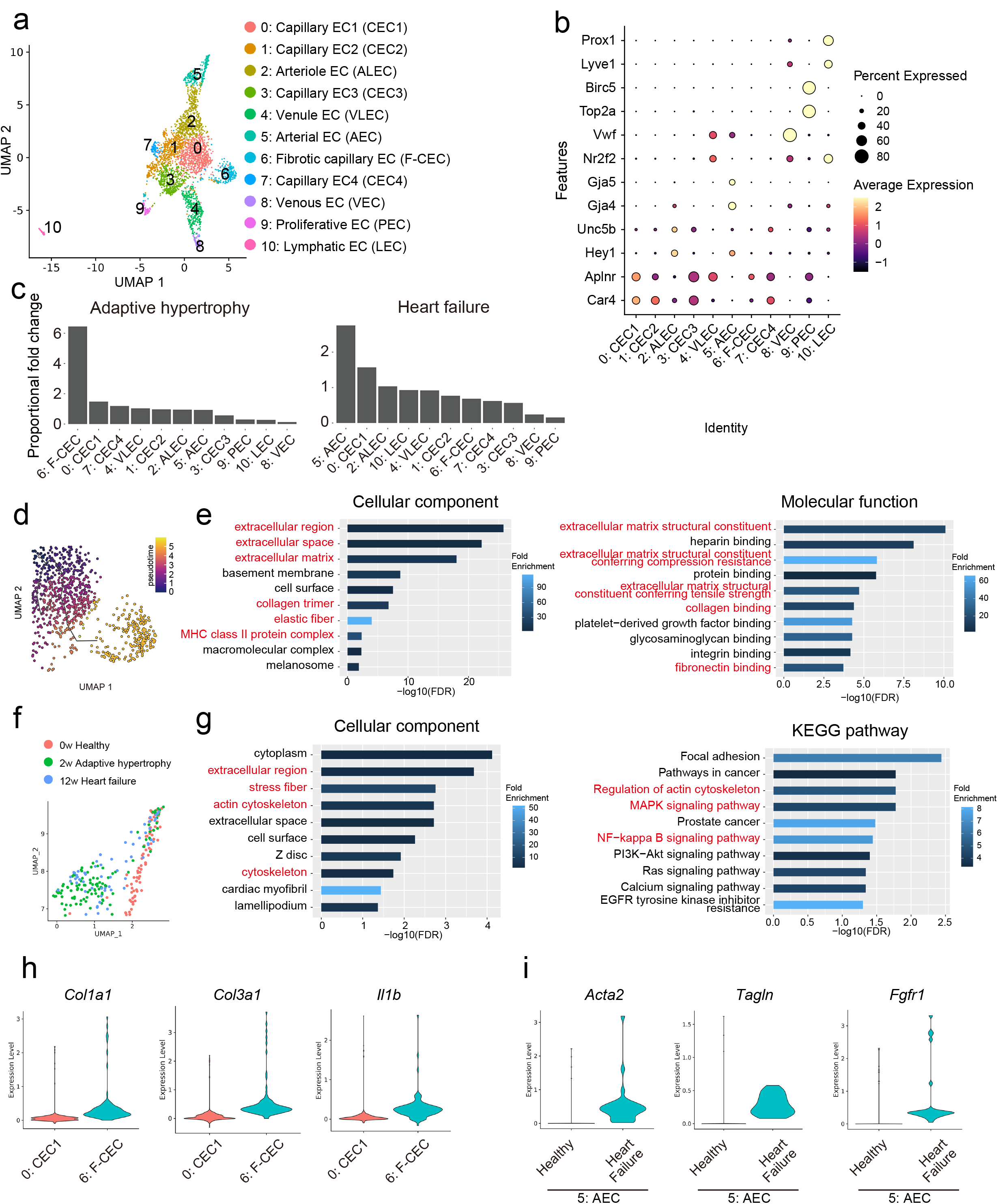
scRNA-seq revealed distinctive endothelial cell subgroup and qualitative change of arterial endothelial cells a, UMAP plot for ECs in combined datasets of healthy, adaptively hypertrophic, and failed hearts. The cells were reclustered before plotting them. **b,** Visualization of EC subgroups and their specific marker genes. Larger dots indicate more cells in the subgroups expressing the marker genes. A brighter colour indicates a higher average expression. **c,** Proportional fold-changes of each subgroup cell number in adaptive hypertrophy and heart failure compared to the healthy state. The subgroups were arranged in descending order of fold-change. **d,** Pseudotime map around subgroup 0 and 6. Brighter colour in pseudotime map indicate more differentiated cells. **e,** GO analysis graphs (left: cellular component; right: molecular function). The rows in the GO graphs were top-10 significant GO terms. X axis was −log10(FDR). Brighter bars indicate higher fold enrichment. **f,** UMAP in subgroup 5. In the UMAP plot, pink, green, and blue dots indicate healthy, adaptive hypertrophy, and failed heart cells, respectively. **g,** GO cellular component graph (left) and KEGG pathway graph (right); **h,** Visualization of the gene expression difference between subgroups 0 and 6 by violin plots. **i,** Visualization of the gene expression difference in subgroup 5 cells in healthy and heart failure phases using violin plots. UMAP, uniform manifold approximation and projection; EC, endothelial cell; GO, gene ontology analysis; KEGG, Kyoto Encyclopedia of Genes and Genomes; EndoMT, endothelial-mesenchymal transition.

### Endothelial-fibroblast interaction through heart failure progression

Next, we performed cell-cell interaction analysis using ligand-receptor pairs^40^. First, the absolute number of interactions for each cell type was calculated. We found that the interactions were most active in the following order: FB, EC, NC, SMC, and BC (Fig. 4a). Next, we analysed which interactions were most active between cell types and revealed that the interaction between ECs as receptors and FBs as ligands was the most prominent (Fig. 4b). Among the receptors in ECs, genes associated with the Scarb family, including *Scarb1* and *Scarb3* (*CD36*), were frequently detected (Fig. 4d). When the total number of interactions in the Scarb family was calculated, FBs were found to be the main source of ligands, similar to the total number of interactions (Fig. 4e). The EC-FB interaction is activated in healthy hearts and contributes to the maintenance of organ homeostasis. However, the total number of EC-FB interactions increased with the progression of heart failure (Fig. 4c), suggesting that it may also have an impact on pathological changes in heart failure development. We calculated the EC-FB interaction, which increased with the progression of heart failure, and clarified that the interaction through the SCARB1 receptor ^41^ was a top-2 interaction (Fig. 4f-h).

**Fig. 4:**
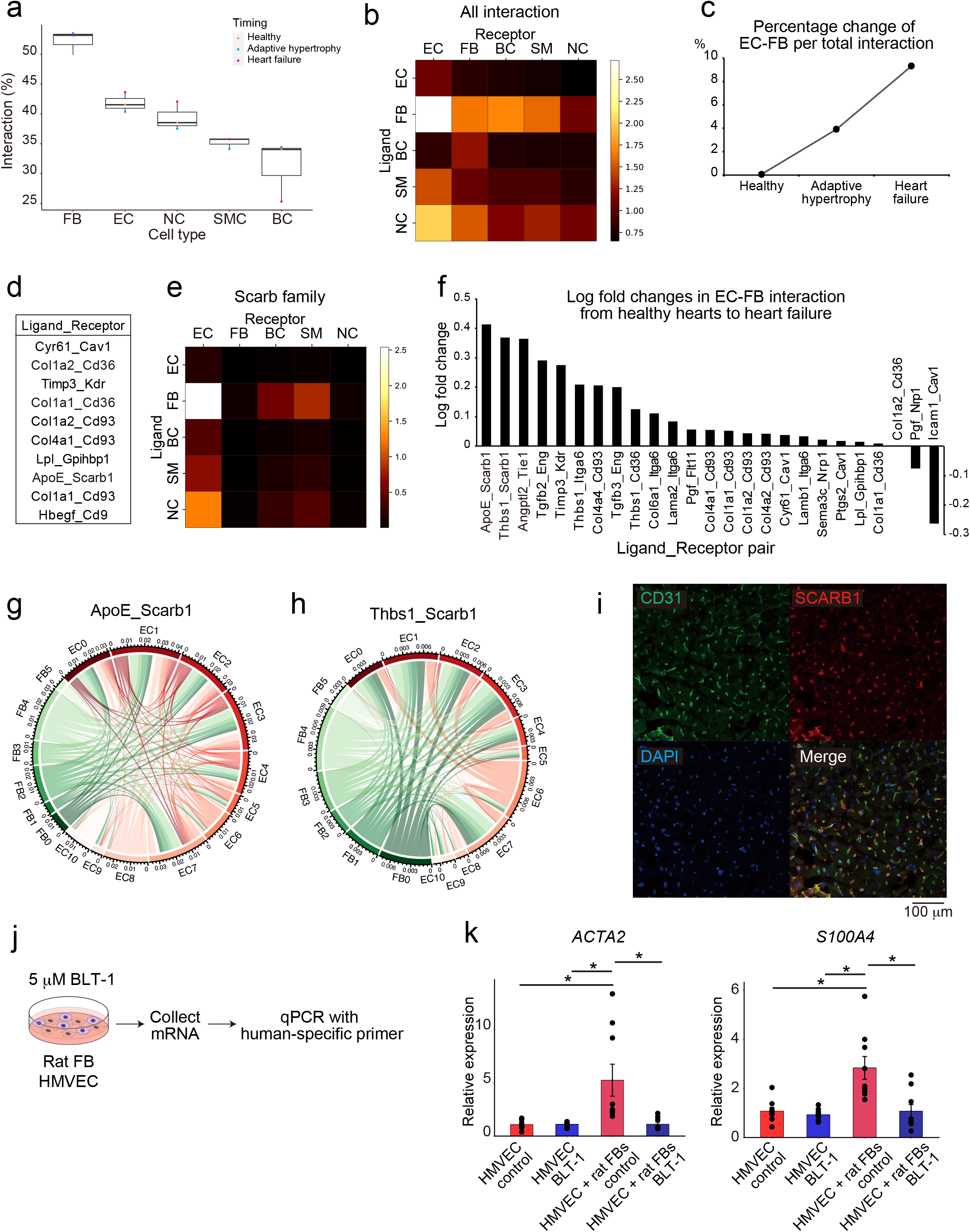
Interaction analysis in scRNAseq analysis and EC-FB co-culture system a, Total number of interactions by cell type. The y-axis shows the total interaction proportion. Variances in healthy, adaptive hypertrophy, and heart failure are shown in the boxplots. **b,** Heatmap of the interactions. The rows and columns represent ligand-expressing and receptor-expressing cell types, respectively. A brighter colour indicates stronger interactions. **c,** Temporal trend of EC-FB interaction in the healthy, adaptive hypertrophic, and heart failure phases. The yaxis represents the percentage change in the total endothelial cell–fibroblast interaction. **d,** The top-10 strongest interaction pairs between ECs and FBs. **e,** Heatmap of scavenger receptor family interactions. A brighter colour indicates stronger interactions between the two groups. **f,** Log-fold changes in EC-FB interaction from the healthy to heart failure phase. The x-axis and y-axis represent the ligand-receptor pairs and log-fold changes in EC-FB interactions in failed hearts, respectively. **g,h,** Circos plots of Scarb1 specific interaction pairs. The outermost layer bandwidth represents the total amount of interactions provided and received. The green and red bands indicate FBs and ECs, respectively. Arcs between bands represent interactions between cell types. The colour and width of the arc correspond to the ligand-expressing cell types and number of interactions, respectively. (g) ApoE-Scarb1 and (h) Thbs1-Scarb1 interactions are displayed. **i,** Representative immunohistochemistry (IHC) images of CD31 (green) and SCARB1(red) in the healthy mouse hearts. Nuclei stained with DAPI (blue). **j,** Experimental schema of EC-FB co-culture. **k,** Results of qPCR for the EC-FB co-culture. *Gapdh* was used as the internal control. Error bars represent standard errors. * indicates p < 0.05 and ** indicates p < 0.01. SM, smooth muscle cells; BC, blood cells; EC, endothelial cells; FB, fibroblasts; NC, neural cells; mRNA, messenger ribonucleic acid; qPCR, quantitative polymerase chain reaction; HMVEC, human cardiac microvascular endothelial cells.

Next, we confirmed the expression of Scarb1 in capillary ECs in the heart by immunofluorescence (Fig. 4i). To investigate whether the signalling pathway mediated by the Scarb1 receptor in ECs, with FBs as ligands, is involved in the pathological changes of ECs in heart failure, we performed co-culture of FBs and ECs (Fig.4j and Extended data Fig.4a). Co-culture of rat fibroblasts and Human Cardiac Microvascular Endothelial Cells (HMVEC) resulted in an increased expression of pathological fibrotic markers such as *ACTA2* and *S100A4* in HMVECs (Fig.4k). This suggests that ECs may acquire characteristics of mesenchymal cells, as observed in our single-cell analysis. Notably, treatment with BLT-1, a specific inhibitor of SCARB1, resulted in the decreased expression of these markers (Fig.4k). This suggests that the initiation of EndMT is mediated by EC-FB crosstalk via SCARB1 in ECs.

### Scarb1 inhibition prevented heart failure progression

To examine whether Scarb1 inhibition improves pathological changes in mouse heart failure models, we administered the SCARB1 inhibitor BLT-1 to TAC-induced heart failure mice (Fig.5a). Cardiac echocardiography revealed that continuous daily administration of BLT-1 from the first day of TAC significantly improved left ventricular ejection fraction (LVEF) dysfunction compared to that in control TAC mice (Fig.5b-d, Extended Data Fig.5a, b). Furthermore, we investigated whether BLT-1 administration reduced cardiac fibrosis, as observed in the co-culture system. Azan staining revealed that perivascular fibrosis was significantly inhibited by BLT-1 treatment (Fig. 5e, f, and Extended Data Fig.5a). Additionally, qPCR showed that the fibrotic markers *Col1a1*, *Tgfb*, and *Timp1*, which were increased by TAC, were decreased to levels comparable to those in the sham group, which did not undergo TAC after BLT-1 administration (Fig. 5g and Extended Data Fig.5c). Moreover, we found that *Postn*, a marker of pathological fibroblasts, was reduced by BLT-1 administration (Fig.5g), indicating the inhibition of overall cardiac fibrosis. We also examined changes in inflammatory responses during the progression of heart failure. Inflammatory molecules, such as *Il1b* and *Il6* were downregulated following BLT-1 administration (Fig.5h). These results suggest that Scarb1 inhibition exerts protective effects by reducing cardiac dysfunction, fibrosis, and the inflammatory response induced by TAC.

**Fig. 5:**
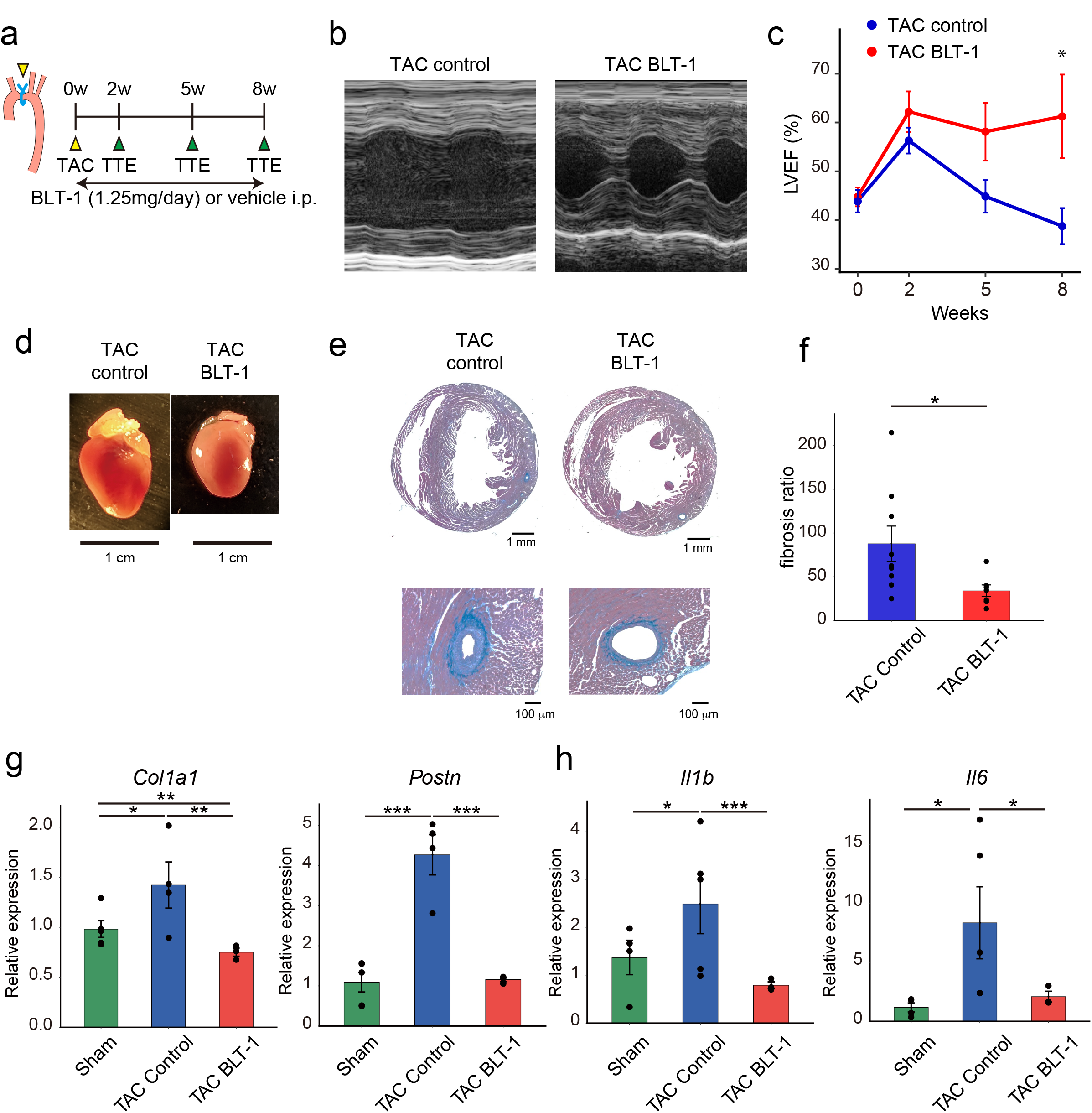
SCARB-1 receptor inhibitor attenuated pressure overload-induced heart failure progression a, Experimental schematic of BLT-1 treatment with TAC. Mice subjected to TAC were randomized into BLT-1 treatment and control groups. From the day of surgery day BLT-1 1.25 mg/body/day or phosphate-buffered saline (PBS) was intraperitoneally injected into the treatment and control groups, respectively. Echocardiography was performed before surgery and at 2, 5, and 8 weeks after surgery. **b,** Representative M-mode echocardiography images of control mice (left) and treatment mice (right). **c,** left ventricular ejection fraction (LVEF) changes over time. The x-axis was post-operative weeks. * indicates p < 0.05. **d,** Representative photos of sacrificed murine hearts. Control (left) and treatment groups (right). **e,** Representative microscopic images of azan-stained sacrificed hearts. Control group x 10 overall picture (upper left), treatment group x10 overall picture (upper right), control group x 20 perivascular picture (lower left), and treatment group x 20 perivascular picture (lower right). **f,** Results of fibrosis ratio quantification in the control and treatment groups (*n* = 3 vs. 3). The fibrosis ratio was calculated as the perivascular fibrosis area divided by the target vessel area. * indicates p < 0.05. **g,** Result of qPCR for the sham, TAC control, and TAC BLT-1 groups (*n* = 4 vs. 5 vs. 3). Two technical replicates were 2 for each measurement, and the values were averaged. *Gapdh* was used as the internal control. Each point represents a single measurement. Error bars represent standard errors. * indicates p < 0.05, ** means p < 0.01, and *** indicates p < 0.001. BLT-1, block lipid transport-1; TAC, transverse aortic constriction.

Next, we investigated whether HDL was involved in the observed phenomena, because SCRAB1 was known as a specific HDL uptake receptor ^42^. We used Dil-HDL to visualize HDL uptake into the tissues (Extended Data Fig. 6a, b). In the liver, there was no significant difference in HDL uptake between the BLT-1 administration and control groups (Extended Data Fig. 6c), suggesting that HDL uptake was not affected by the BLT-1 dose used in this study. Additionally, we confirmed that HDL uptake did not occur in the heart regardless of BLT-1 administration (Extended Data Fig. 6d). Therefore, the cardioprotective effect of BLT-1 may be mediated by direct signalling through Scarb1, rather than an action mediated by HDL.

### Scarb1 functions in endothelial cells

To analyse how SCARB1 in ECs is involved in the progression of heart failure, we administered BLT-1 to HMVECs and performed RNA sequencing analysis. Principal component analysis and clustering revealed distinct gene expression clusters in the control and BLT-1 groups (Fig.6a and Extended Data Fig.7a). We conducted a Gene Set Enrichment Analysis (GSEA) and found a significant decrease in gene groups associated with epithelial-mesenchymal transition (EMT) upon BLT-1 administration (Fig.6b), which is similar with the gene groups of EndMT ^43,44^. Therefore, the Scarb1 receptor may directly regulate genes related to fibrosis, as observed in the scRNA-seq analysis and BLT-1 administration in mice. Indeed, as shown in the heatmap, the expression of *ACTA2*, *TAGLN*, and *COL1A1* decreased following BLT-1 administration (Fig.6c), as well as in the scRNA-seq analysis, suggesting that Scarb1 inhibition could serve as a novel therapeutic approach to suppress fibrosis. The heatmap also showed *ID1*, *ID2*, and *ID3* expression, which are downstream genes in the BMP signalling pathway, was decreased by BLT-1 (Fig.6c). Furthermore, we observed that pre-inflammatory molecules such as *IL1B* and *VCAM1* and chemokines such as *CCL2*, *CXCL1* and *CXCL12*, were decreased by BLT-1 treatment (Fig. 6c), indicating the potential involvement of SCARB1 in the inflammatory signalling pathway.

**Figure 6:**
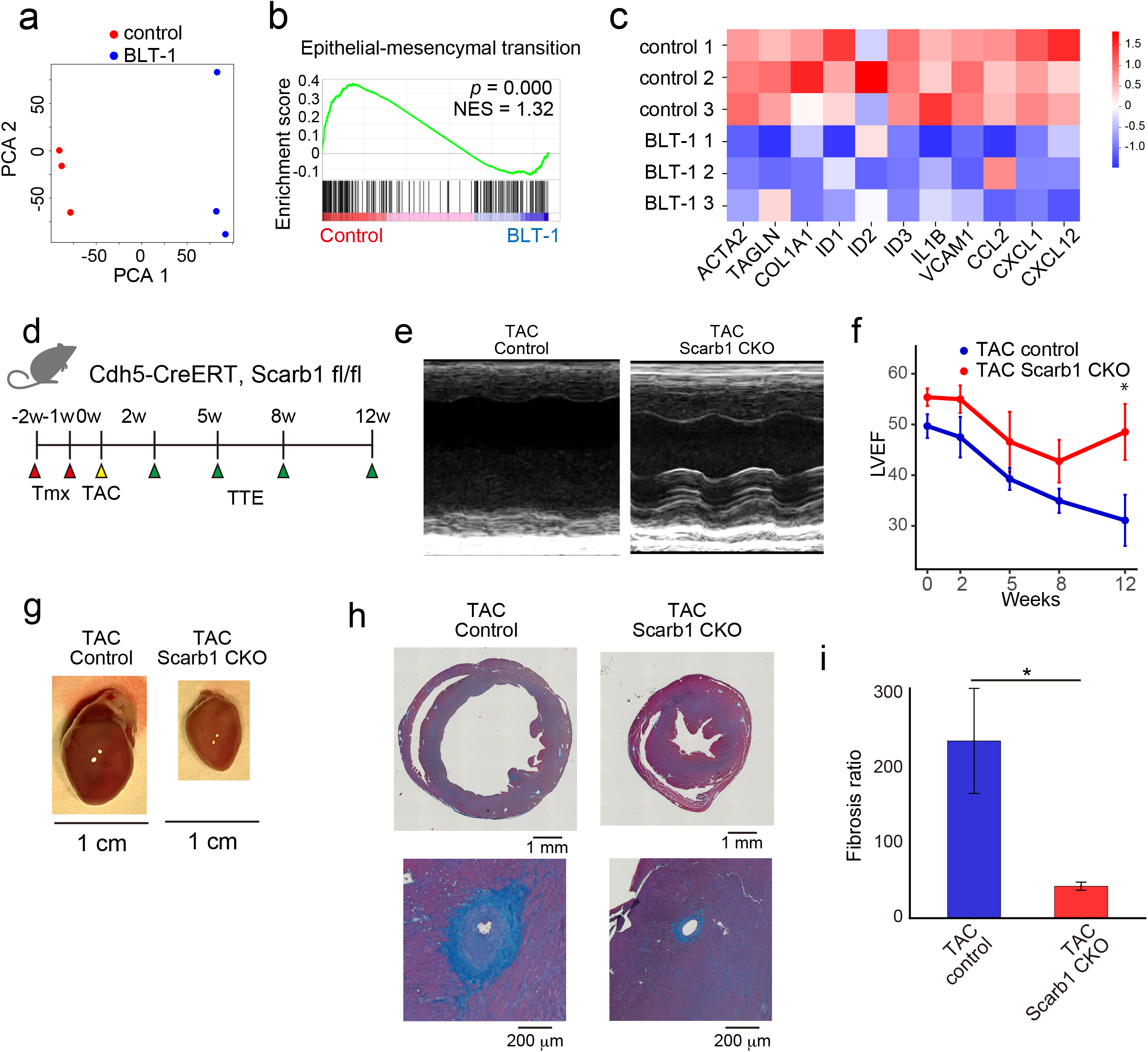
EC-specific Scarb1 knockout ameliorated heart failure condition in pressure-overloaded heart a, Principal component analysis (PCA) of RNA sequences for control and BLT-1 groups (*n* = 3). HMVEC were treated with DMSO (control) and 1 μM BLT-1 for 24 h. **b,** Characteristic results of gene set enrichment analysis (GSEA) of RNA sequences. The green line indicates the enrichment score. **c,** Heatmap of representative differentially expressed genes between the control and BLT-1 treatment groups. Rows represent samples, and columns represent representative genes. Values were Z-score. **d,** Experimental scheme for EC-specific Scarb1 conditional knockout in TAC mice. Before surgery, at weeks 2 and 1, tamoxifen was intraperitoneally injected into (Cdh5-CreERT, Scarb1 fl/fl) mice, and (Cdh5-CreERT, Scarb1 +/+) mice. Echocardiography was performed before surgery and at 2, 5, 8, and 12 weeks after surgery. **e,** Representative M-mode echocardiography images of control (left) and Scarb1 CKO mice (right). **f,** in left ventricular ejection fraction (LVEF) change over time. The x-axis was post-operative weeks. * indicates p < 0.05. **g,** Representative photographs of sacrificed murine hearts. Control (left) and treatment groups (right). **h,** Representative microscopic images of azan-stained sacrificed hearts. Control group x 10 overall picture (upper left), treatment group x 10 overall picture (upper right), control group x 20 perivascular picture (lower left), and treatment group x 20 perivascular picture (lower right). **i,** Quantification of the fibrosis ratio in the control and treatment groups (*n* = 9 vs. 6). The fibrosis ratio was calculated as the perivascular fibrosis area divided by the target vessel area. * indicates p < 0.05.

Finally, to investigate the EC-specific functions of *Scarb1* in the progression of heart failure, we generated EC-specific Scarb1 knockout mice (Scarb1 CKO) by crossing Cdh5-CreERT2 mice expressing endothelial cell-specific Cre induced by tamoxifen with *Scarb1*-flox mice (Fig. 6d). qPCR analysis confirmed the downregulation of *Scarb1* expression in endothelial cells after tamoxifen administration (Extended Data Fig.7b). Heart failure was induced by performing TAC after tamoxifen administration in *Scarb1* CKO and control mice (Fig. 6d). Cardiac echocardiography revealed that the impairment of cardiac function induced by TAC was rescued in *Scarb1* CKO mice (Fig. 6e-g and Extended Data Fig.8a). Additionally, vascular fibrosis was suppressed in *Scarb1* CKO mice (Fig. 6h, i; Extended Data Fig.8b).

## Discussion

In this study, we clarified that the EC-FB interaction through *Scarb1* is important for pathological changes in ECs during the development of heart failure. Both capillary and arterial ECs acquired mesenchymal characteristics at the heart failure stage according to scRNA-seq analysis, indicating that EndMT plays an important role in the pathological changes in heart failure. Notably, co-culture of ECs and FBs revealed that FBs directly interacted with ECs, which have mesenchymal characteristics, and SCARB1 inhibition reduced these changes. Moreover, both SCARB1 inhibition and EC-specific *Scarb1* knockout rescued TAC-induced heart failure and cardiac fibrosis, suggesting *Scarb1* in ECs plays a key role in the induction of worsening heart failure and is a potential therapeutic target.

A previous report showed that TAC-induced heart failure was exacerbated in systemic *Scarb1* -/- mice, and that hepatocyte-specific gene transfer of *Scarb1* rescued heart failure^45^, contradicting our results. However, we believe that these two results are consistent for several reasons. Ilayaraja et al. concluded that the hepatocyte-specific function of *Scarb1* is important for maintaining cardiac function because cardiac dysfunction induced in *Scarb1* -/- mice was rescued by AdSR-BI gene transfer to liver cells^45^. In contrast, our mouse model used tamoxifen-induced EC-specific *Scarb1* knockout mice, and we showed that the EC-specific deletion of *Scarb1* did not affect HDL metabolism in the liver. A previous report showed that EC-specific *Scarb1* knockout mice reduced atherosclerosis progression^46^, which is similar to our cardioprotective results. Therefore, we hypothesized that *Scarb1* in ECs exacerbates heart failure by showing fibrotic tendencies when stimulated by FBs, whereas *Scarb1* in the liver is cardioprotective and normalizes HDL metabolism. Although BLT-1 may inhibit systemic SCARB1, our data did not show an effect on HDL uptake in the liver, suggesting that BLT-1 primarily acts on ECs to exert its cardioprotective effects.

Although several pathological changes such as decreased NO production, cellular senescence, and inflammation occur in ECs through organ dysfunction, our scRNA-seq analysis showed that the most noticeable change was the acquisition of mesenchymal characteristics in both capillary and arterial ECs. Moreover, we revealed that FBs induced these changes in ECs via SCARB1 and that SCARB1 blockage in ECs reduced mesenchymal changes. These results suggest that EndMT via SCARB1 signalling is important for the progression of hypertensive heart failure. EndMT is involved in the progression of heart failure in various aspects such as cardiac fibrosis ^38,39^. However, the detailed mechanism through which EndMT exacerbates heart failure remains unknown. Further studies are necessary to clarify the downstream mechanisms of SCARB1 involved in EndMT in the next step.

Our results suggest that ECs are potential therapeutic targets for heart failure. ECs play a crucial role in the progression of hypertensive heart failure, which was also examined in this study, particularly in the transition from the adaptive hypertrophy phase to the noncompensatory heart failure phase, when cardiac function irreversibly declines. Although neither BLT-1 administration and *Scarb1*-CKO mice with TAC treatment prevented adaptive hypertrophy (Extended Data Fig. 5b and 8a), their cardiac function was preserved (Fig. 5c and 6f). This suggests that inhibiting *Scarb1* in EC could prevent the transition from adaptive hypertrophy to heart failure. Hypertensive heart failure is the most common cause of heart failure in Japan. Thus, we believe that our study will play an important role in the future development of heart failure treatments that target pathological endothelial cells.

Although we revealed important findings in this study, there are several limitations. RNA-seq analysis showed *ID1*, *ID2* and *ID3* were downregulrated by BLT1 administration (Fig. 6c), indicating BMP signalling, which is involved in cardiovascular disease and cardiac fibrosis^47,48^, has an important role in pathological EC changes. However, detailed mechanism of the downstream signals of Scarb1 cannot be fully understood in this study. Furthermore, the ligand of SCARB1 in the EC-FB interaction has not been elucidated. In the future, it will be necessary to conduct a detailed analysis of how Scarb1 is involved in EndMT, including the identification of ligands and downstream pathways. In this study, we have focused on non-cardiomyocytes, and cardiomyocytes were not included in analysis. The relationship between Scarb1 signal and cardiomyocytes should be clarified in next study. Further research based on the findings from this study may yield important findings for future clinical applications.

## Data availability

The scRNA-seq and RNA-seq data will be publicly available after formal publication of the paper.

## Acknowledgements

We thank all the members of our laboratory for their assistance. This research was supported by AMED under Grant Number 22ek0210169h0001, Bayer Academic Support, MSD Life Science Foundation, Japanese Heart Failure Society, and Takeda Science Foundation.

## Contributions

D. K. designed the experiments. T.K., M.K., Y.A., J.K., T.N., T. K., and H.H. collected data. T.K. analysed the data. Y.S. conducted scRNA-seq analysis. Y.K. provided experimental materials. S.Y.,K.F. and M.I. supervised the study. D.K. and T.K. prepared the manuscript.

## Competing interests

K.F. is a Founding Scientist funded by the SAB of Heartseed Co., Ltd. D.K., T.K., M.K., Y.A., J.K., T.N., Y.S. and S.Y. declare no competing interests.

## Methods

### Animals

The animal procedures followed the Guide for the Care and Use of Laboratory Animals published by the US National Institutes of Health (Publication no. 85-23, revised 1996), and the study protocol was approved by the Institutional Animal Care and Use Committee of Keio University School of Medicine. *Scarb1* flox mice (C57BL/6J background) were provided by Dr. Thierry Huby (Sorbonne Université). *Cdh5*-CreERT2 mice were purchased from the Center for Animal Resources and Development, Kumamoto University. These mice were originally developed by Dr. Kubota (Keio University Hospital) ^31^. All mice were housed in climate-controlled (23 °C) specific pathogen-free facilities with a 12-h light-dark cycle and free access to standard laboratory food (CE2; CLEA Japan Inc.) and water at Keio University.

### Transverse aortic constriction (TAC)

Inhalation anaesthesia with 4% isoflurane (Fujifilm, Tokyo, Japan) was administered to eight-week-old male C57BL/6J JCL mice (Nihon CREA, Tokyo, Japan). After the mice were sedated, 1% isoflurane was administered. Twenty-two-gauge outer cannula (Terumo, Tokyo, Japan) was inserted into the trachea. A respirator was connected to the cannula and artificial breathing was established by 0.1 mL × 120 /min. The mice were placed in the supine position and their chest hair was removed using an epilation cream (Kracie, Tokyo, Japan). The chest walls were gently washed and sterilized with 70% ethanol (Kaneichi, Osaka, Japan). Approximately 2 cm midline skin incision was made on the chest. The second rib was opened and the thymus was pushed towards the head. Fat and connective tissues were carefully detached from the aortic arch. The arch was ligated using a 10-0 nylon thread (Natsume, Tokyo, Japan) with 29-gauge needle (Nipro, Osaka, 50. Japan) spacer. Two knots were made and atrial enlargement was observed. The spacer was removed and blood flow resumption was confirmed by atrial contraction. The opened ribs and skin were sutured using 5-0 silk thread (Natsume, Tokyo, Japan). Anesthesia was suspended, and intubation was removed after the recovery of spontaneous breathing. The mice were allowed to recover on a 37 °C heat pad.

### Echocardiography

Inhalation anaesthesia was introduced and the hair on the chest wall was removed, similar to the TAC procedure. First, parasternal long axis view was obtained, and the optimal echo probe (Veve 2100, FujiFilm VisualSonics, Tokyo, Japan) position was adjusted. Next, the probe was rotated just above the papillary muscles and a short-axis view was obtained. An M-mode video was recorded, and the interventricular septum thickness (IVS), posterior left ventricular wall thickness (PWT), left ventricular end-diastolic dimension (LVDd) and left ventricular internal dimension in systole (LVDs) were measured. The left ventricular ejection fraction was automatically calculated from LVDd and LVDs. Echocardiography was performed at 0, 2, 5, and 8 weeks after TAC in BLT-1 administration mice, and at 0, 2, 5, 8, and 12 weeks after TAC in Scarb1 CKO mice.

### Preparation of single-cell suspension

Mice were euthanized using the cervical dislocation technique. The rib cage was quickly opened with scissors and 10 mL of ice-cold phosphate-buffered saline (PBS) was intermittently injected into the right ventricle after the left atrium incision. When the lungs turned white, the hearts were extracted and soaked in ice-cold PBS. The atria were removed in cold PBS. Type II collagenase (Worthington, CA, USA) (13.5 g) and DNase I (2 mg (Qiagen) were dissolved in 30 mL of

F12Ham medium (Thermo Fisher Scientific, MA, USA). The heart was minced using scissors in an ice-cold Petri dish (Corning, NY, USA) and mixed with a collagenase solution. The solution was poured into an Erlenmeyer flask soaked in a 38 °C warm bath (Taitec, Saitama, Japan) with a magnetic stirrer (AS ONE, Osaka, Japan) for 30 min. After a 30-min incubation, the mixture was repeatedly cannulated using an 18G needle (Nipro, Osaka, Japan) with a 10 mL syringe (Terumo, Tokyo, Japan). After cannulation, a 30-minute reaction was performed. Ice-cold PBS (20 mL PBS to stop the reaction. This solution was centrifuged at 450 × *g* and 4 ℃. The supernatant was removed, and 5 mL of 1X RBC Lysis Buffer (Thermo Fisher Scientific, MA, USA) was added on ice with gentle shaking. Ice cold PBS (45 mL) was added and the mixture was passed through 70 µm and 40 µm filter (pluriSelect, Leipzig, Germany) consecutively. The solution was centrifuged as previously described and the supernatant was removed. CellCover (3 mL Anacyte Laboratories, Hamburg, Germany) was added on ice for 5 min. Ice-cold PBS (47 mL) was added and centrifuged as described previously. The supernatant was removed, and 4 mL of ice-cold PBS was added. In the 15 mL tube (Corning, NY, USA), 3 mL “bottom Percoll,” 4 mL “top Percoll,” and the previous solution of 1 mL was stacked in this order. The “bottom Percoll” and “top Percoll” comprised the following:

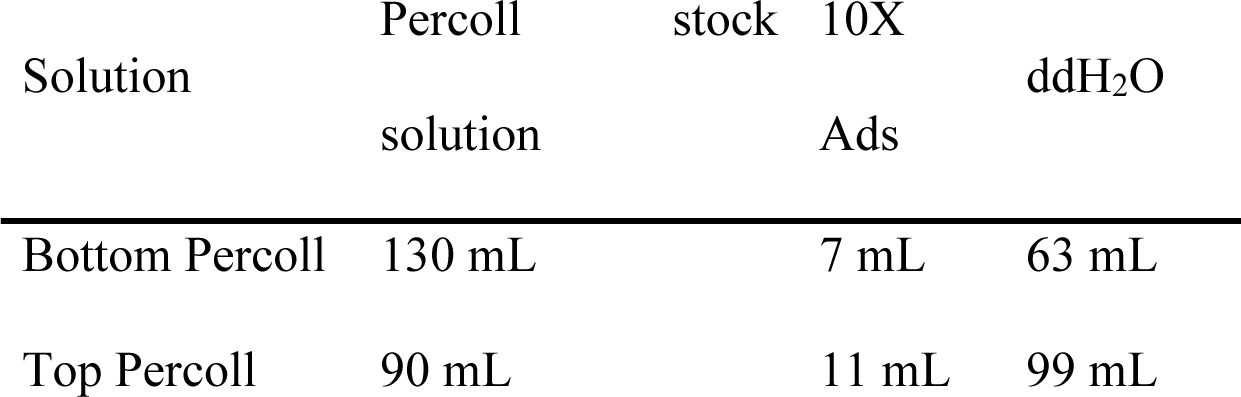

A Percoll stock solution was prepared by mixing 450 mL of Percoll (Sigma-Aldrich, Darmstadt, Germany) with 50 mL of 10 X Ads. The ingredients of the 10 X Ads were as follows:

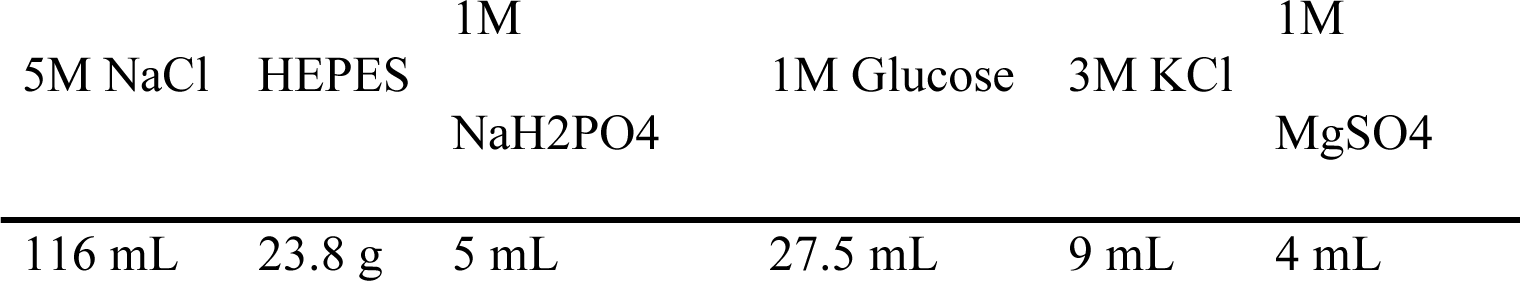

The stacked solution was centrifuged for 25 minutes at 3000 × *g* (Beckman, CA, USA) and 25 ℃. The debris layer and top Percoll were swiftly aspirated. The cell layer and bottom Percoll were moved to a new tube and diluted with ice-cold PBS up to 50 mL and centrifuged for 6 min at 450 × *g* and 4 ℃. The supernatant was removed, and the precipitate was washed in the same manner. Finally, 1mL of CellCover was added before application to the single-cell analysis apparatus.

### Single-cell RNA sequencing, mapping and exploratory analysis

This solution was applied to a Chromium Controller (10X Genomics, CA, USA) using a manufacturer-defined protocol^28^. The RNA solution was then applied to the HiSeq 3000 system (Illumina, CA, USA). Read count data were processed using Cell Ranger (10X Genomics, CA, USA) and analysable UMI count data were retrieved.

To reproducibly analyze the sequence data, we used R 4.2.1, Seurat 4.1.0, Monocle 3 v1.3.1, and circlize 0.4.15 packages. To convert UMI count data into analysable “Seurat object,” “Read10X” and “CreateSeuratObject” functions in Seurat package were used. The options for the latter were minimum cell = 3 and minimum feature = 200. “NormalizeData” function in Seurat package was used for data normalization. To each dataset (TAC 0 W, 2 W, 12 W) “FindVariableFeatures” function in Seurat package was applied to extract key genes for clustering cells that are called “features.” The options were selection.method = “vst” and nfeatures = 2000. To remove the batch effect Seurat package employs “anchor” approach that uses metric learning. “SelectIntegrationFeatures” function selected key genes for “anchor” cells that were utilized for connecting nodes of different datasets. “FindIntegrationAnchors” function used these genes to identify anchor cells. Finally, the “IntegrateData” function integrated the three different timing datasets into one. The integrated datasets were scaled, dimensionally reduced and clustered by “ScaleData,” “RunPCA,” “RunUMAP,” “FindNeighbors” and “FindClusters” functions. Options were npcs = 30, reduction = “pca,” dims = 1:30 and resolution = 0.5. These functions also employ metric learning methods, including the k-nearest neighbour and shared nearest-neighbour graph methods. After the first clustering, crude cell type populations (endothelial cells, smooth muscle cells, fibroblasts, blood cells and neural cells) were identified by known cell-type specific markers (“Pecam1,” “Cdh5,” “Kdr,” “Myh11,” “Tagln,” “Acta2,” “Pdgfra,” “Col1a1,” “Tcf21,” “Ptprc,” “Itgam,” “Kcna1,” “Kcna2” and “Sox10”). For further investigation of the subtypes, we divided the combined dataset into subdatasets of each crude cell type and re-clustered them using the same settings. Re-annotation was performed using the genes shown in the figures. Specifically endothelial cells were annotated by “Fabp4”, “Fabp5”, “Car4”, “Aplnr”, “Dll4”, “Hey1”, “Unc5b”, “Gja4”, “Gja5”, “Nr2f2”, “Vwf”, “Top2a”, “Birc5”, “Lyve1” and “Prox1”. The Chiefly Monocle 3 package was used to visualize local gene expression transitions and cell trajectories. Seurat data objects were converted into “cell_data_set” objects. They were clustered, embedded in graphs, and aligned using the Monocle 3 function as a vignette showed (https://github.com/satijalab/seurat-wrappers).

### Gene ontology analysis

To identify key genes that discriminate certain populations from others, we employed “FindMarkers” function in Seurat package. The options were logfc.threshold = 0.1, and min.pct =

0.1. The output genes were screened using a threshold of p < 0.05, and the top 150 genes were cast into the Database for Annotation, Visualization, and Integrated Discovery (DAVID) webapp. The GO (BP, CC, and MF) and KEGG pathway results were extracted.

### Interaction analysis

To quantify cell-cell interactions, we conducted matrix calculations. We sliced the known ligand and receptor (https://pubmed.ncbi.nlm.nih.gov/28614297/) matrix from the raw UMI count matrix. The expression of each gene was averaged within each sample week (0 W, 2 W and 12 W) or cell-count-weighted averaged through subtypes, depending on the subsequent analysis. These expression matrices were scaled by the sum of the matrices themselves and multiplied by 100000. Given that the ligand column vector was *L* and the receptor row vector was *R*, the adjacent matrix of each cell type (*A*) was calculated as follows:

*A* = *L* x *R* (non-interchangeable)

These adjacent matrices were visualized by circlize package as circos plots.

### Scarb-1 receptor blocker treatment of TAC mice

TAC was performed as previously described. Mice were randomized into treatment or control groups. Fifty microliters of block lipid transport-1 (BLT-1) solution in 1 mL PBS was intraperitoneally injected into the treatment group mice once a day. The BLT-1 solution consisted of 25 mg of BLT-1 and 1 mL of DMSO. Echocardiography was during pre-operation, and at post-operative weeks 2, 5 and 8.

### Quantification of HDL in vivo

To test the effect of BLT-1 on the plasma concentration of high-density lipoprotein (HDL), we collected blood samples from the right atrium of sacrificed mice. SRL Inc. quantified the high-density lipoprotein (HDL) concentrations in these samples.

### Conditional knock-out of *Scarb1* gene in endothelial cells

*Cdh5*-CreERT2/*Scarb1*^flox/flox^ and *Cdh5*-CreERT2/ *Scarb1*^+/+^ mice were generated. Tamoxifen oil was intraperitoneally injected (45 mg/BW kg) into these mice two and one weeks before the TAC operation. Tamoxifen oil was prepared by diluting 50 mg tamoxifen (Sigma-Aldrich, Darmstadt, Germany) in 1 mL ethanol (Wako, Tokyo, Japan) and 9 mL sunflower oil (Sigma-Aldrich, Darmstadt, Germany). TAC was performed as previously described. Echocardiography was performed during pre-operation, and at post-operative weeks 2, 5, 8 and 12.

### Perivascular fibrosis quantification

In the Azan-stained specimen, the fibrosis ratio was calculated as follows: Fibrosis ratio = A_F_/abπ (no unit given A_F_ was the area of fibrosis, a/b was the long/short axis lengths of the target vessel and π was the circular constant.

### RT-PCR

Total RNA was dissolved in TRIZOL (Thermo Fisher Scientific, MA, USA), separated using chloroform (Wako, Tokyo, Japan), precipitated with isopropyl alcohol (Wako, Tokyo, Japan), and purified using 70% ethanol (Wako, Tokyo, Japan). ReverTra Ace qPCR RT Master Mix with gDNA Remover (Toyobo, Osaka, Japan) was used for reverse transcription.

The primers used were as follows.

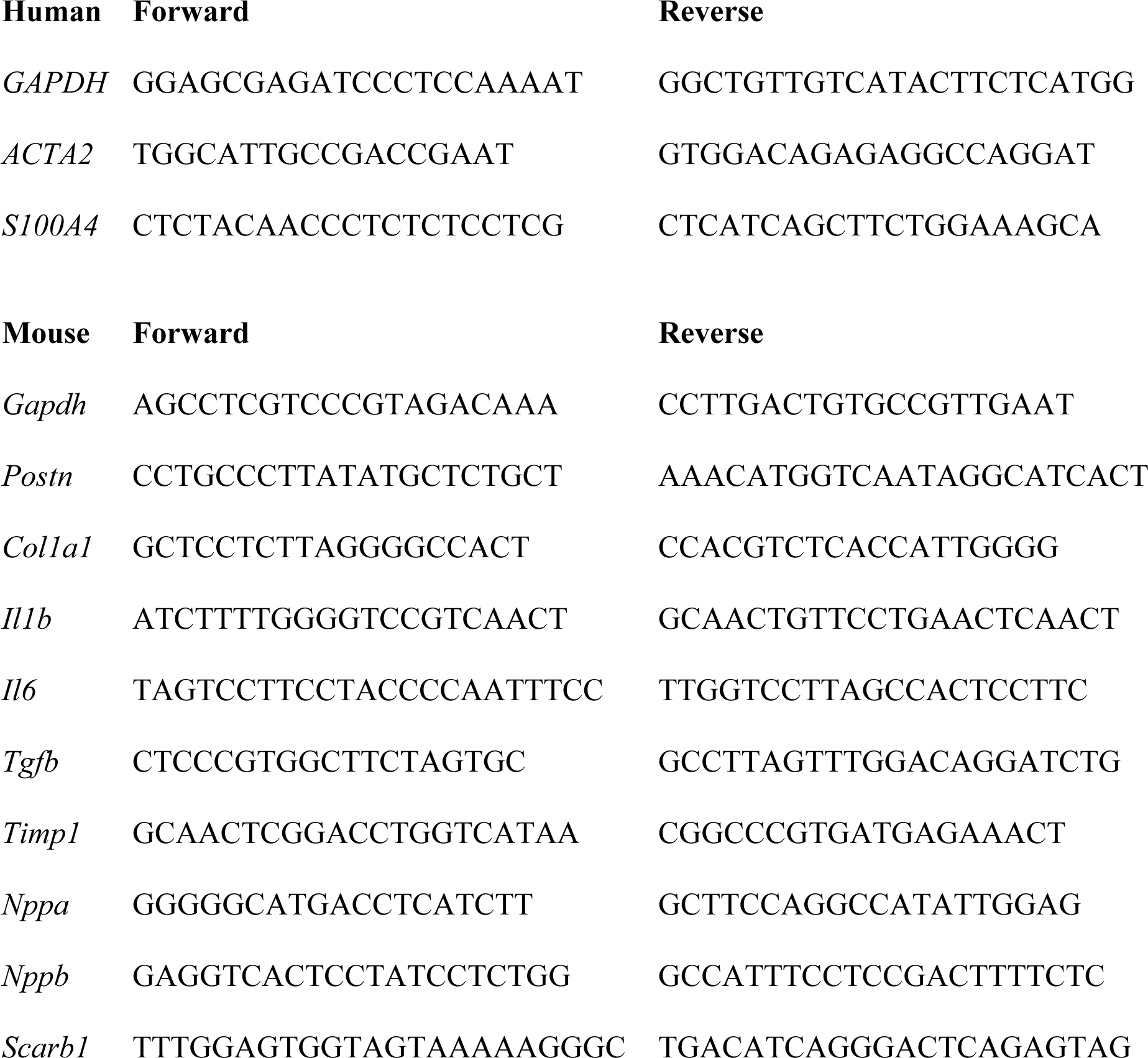

### Co-culture of endothelial cells and fibroblasts

Human cardiac microvascular endothelial cells (HMVEC Cardiac MV Endo, batch number 0000550176, Lonza, Basel, Switzerland) and rat heart fibroblasts separated from 3–7 day-old Jcl:Wistar rats (CLEA Japan Inc., Tokyo, Japan). In the co-culture group, they were seeded together in each well of a 6-well plate (Corning, NY, USA) by 1×10^5^cells/2.5 mL and 0.5×10^5^cells/2.5 mL final concentrations respectively. In the monoculture group, HMVEC were seeded at a final density of 2×10^5^cells/2.5mL final concentration. In each treatment well, a final concentration of 5 µM BLT-1 was added. Dimethyl sulfoxide was added to each control well to ensure that the final DMSO concentration was the same as that in the treatment group. The culture medium used was EGM^TM^-2 MV Microvascular Endothelial Cell Growth Medium-2 BulletKit^TM^ (Lonza, Basel, Switzerland). The cells were cultured for 24 h and total RNA was collected as described in the RT-PCR protocol.

### RNA-sequence of BLT-1 treated HMVEC-C

HMVEC were seeded in a 6-well plate at a density of 1 × 10^5^ cells/2.5 mL concentrations. The medium was the same as that used for the co-culture. After 48 h, 1 µM final concentration BLT-1 was added to treatment wells and DMSO was added so that the final DMSO concentration was the same as the treatment group. There were six cultured treatments and controls. Total RNA was collected as previously described, and samples were combined into 3 vs. 3 for RNA sequencing. RNA was purified using the ReliaPrep RNA Cell Miniprep System (Promega, WI, USA). The sequencing and mapping were performed by Macrogen (Tokyo, Japan). The differentially expressed genes between the two groups (p < 0.05) were subjected to Gene Set Enrichment Analysis (GSEA) using the Broad Institute website (https://www.gsea-msigdb.org/gsea/index.jsp).

### Visualization of HDL dynamics in vivo

To delineate the dynamics of HDL, 30 µg/body 1,1’-dioctadecyl-3,3,3’,3’-tetramethylindocarbocyanine perchlorate labelled HDL (Dil-HDL: KALEN Biomedical, MD, USA) was used. Dil-HDL was retro-orbitally injected into the target mice 6 h before sacrifice. Mice were sedated with 1% isoflurane through this procedure.

### Immunohistochemistry

The sample mice were euthanized, and the PBS-perfused hearts and livers were harvested as previously described. Cryosections of these organs were obtained using Tissue-Tek O.C.T. compound (Sakura Finetek Japan, Tokyo, Japan). The specimens were washed under running tap water and PBS. Antigen retrieval was performed by soaking the samples in citrate buffer (pH 6.0) and boiling them twice in a microwave. Citrate buffer was prepared with 2.95 g trisodium citrate dihydrate (Fujifilm, Tokyo, Japan) and 1 L ultrapure water and titrated with 1 M HCl solution (Fujifilm, Tokyo, Japan). Finally, Tween 20 (0.5 mL) was added (Sigma-Aldrich, Darmstadt, Germany).

Targeting endothelial CD31 and SCARB 1, Mouse/Rat CD31/PECAM-1 antibody (R&D Systems, MN, USA), and SR-BI Antibody - BSA free (Novus Biologicals, CO, USA) were utilized. The primary antibodies were dissolved in Blocking One P (Nacalai Tesque, Kyoto, Japan) to obtain a final concentration of 1:100 v/v. Specimens were incubated at 4 ℃ overnight. The specimens were then washed thrice with PBS.

When SCARB1 was immunostained, signal amplification was performed using TSA PLUS CYANINE 5.5 (Perkin Elmer, MA, USA), according to the manufacturer’s protocol. Briefly, the internal peroxidase activity was quenched by incubation with hydrogen peroxide (Wako, Tokyo, Japan) for 30 min at 25 °C before incubation with the primary antibody. Anti-rabbit HRP (Global Life Sciences Technologies Japan, Tokyo, Japan) incubation was performed for 30 min at 25 °C after the primary antibody incubation. The specimens were washed thrice in PBS and incubated in TSA Plus working solution for 10 min at 25 °C.

Alexa Fluor 488 donkey anti-goat IgG (Life Technologies, CA, USA) and Alexa Fluor 594 chicken anti-rabbit IgG (Life Technologies, CA, USA) were used as secondary antibodies for CD31 and SCARB1, respectively. The secondary antibodies were dissolved in Blocking One P to obtain a final concentration of 1:1000 v/v. The cells were then incubated for 1 h at 25 °C.

The specimens were washed in PBS three times and incubated in 1:2000 v/v DAPI (BD Bioscience, NJ, USA) in PBS solution for 5 min.

