## Supplementary Figure 1 - 8 for "Endothelial-fibroblast interactions during Scarb1 accelerate heart failure"

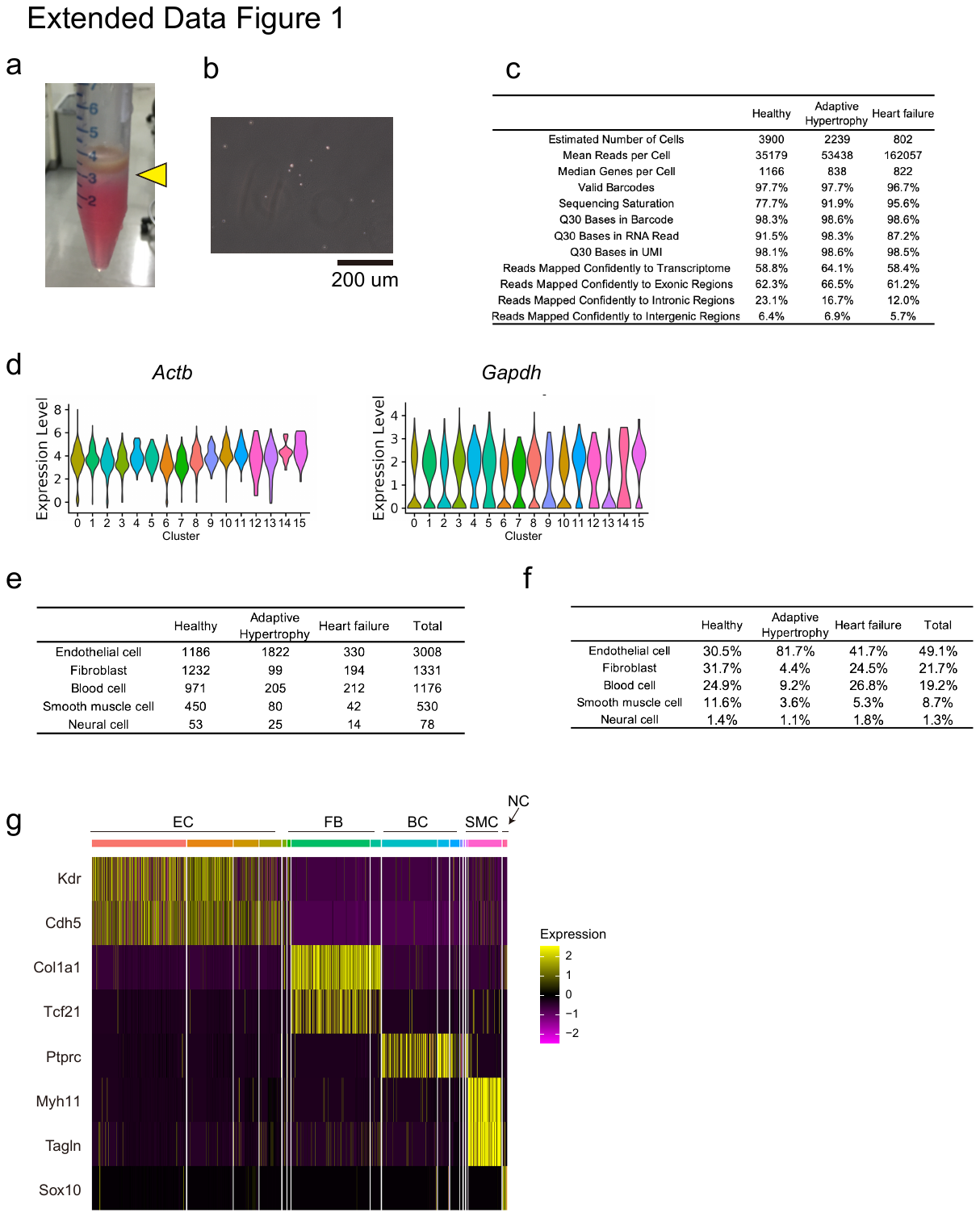

**Extended data fig.1: Acquiring single-cell expression profile of failing hearts.**

a, Murine heart cells were dissociated by collagenase and separated by Percoll. The yellow arrow indicates the boundary where non-cardiomyocytes were concentrated. b, Microscopic photograph of the previous cells. The cells were then resuspended in PBS. c, Summary of scRNA-seq results for healthy, adaptive hypertrophic, and failed heart samples. d, Expression levels of housekeeping genes displayed in violin plots. Actb (left) and Gapdh (right). e-f, Cell numbers annotated in these experiments. Rows were cell types. The columns represent the disease progression phases. Crude cell number (e) and proportion (f). g, Heatmap of characteristic gene expression in each cell type. Rows were genes. Columns represent cell types. Brighter colors indicate stronger expression. PBS, phosphate-buffered saline.

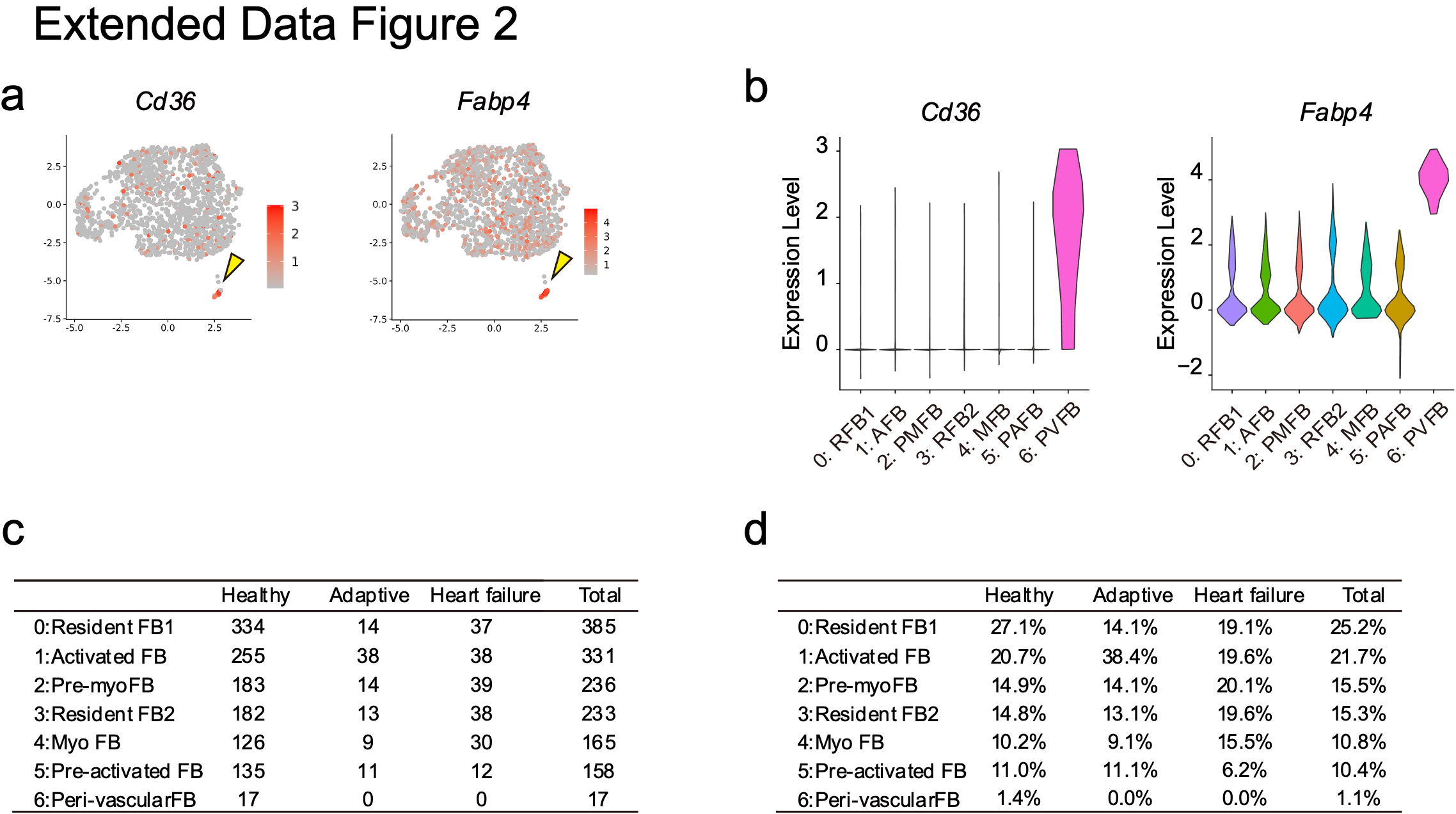

**Extended data fig.2: delineating fibroblasts characteristics in single cell level.**

a-b, Visualization of the PVFB-specific gene expression profile. Featureplots of Cd36 (a, left) and Fabp4 (a, right). Violin plots of the same genes (b). PVFB highly expressed lipid metabolism genes. c-d, Number of annotated cells in these experiments. Rows were cell types. The columns represent the disease progression phases. Crude cell numbers (c) and proportions (d). PVFB, perivascular fibroblasts; FB, fibroblasts.

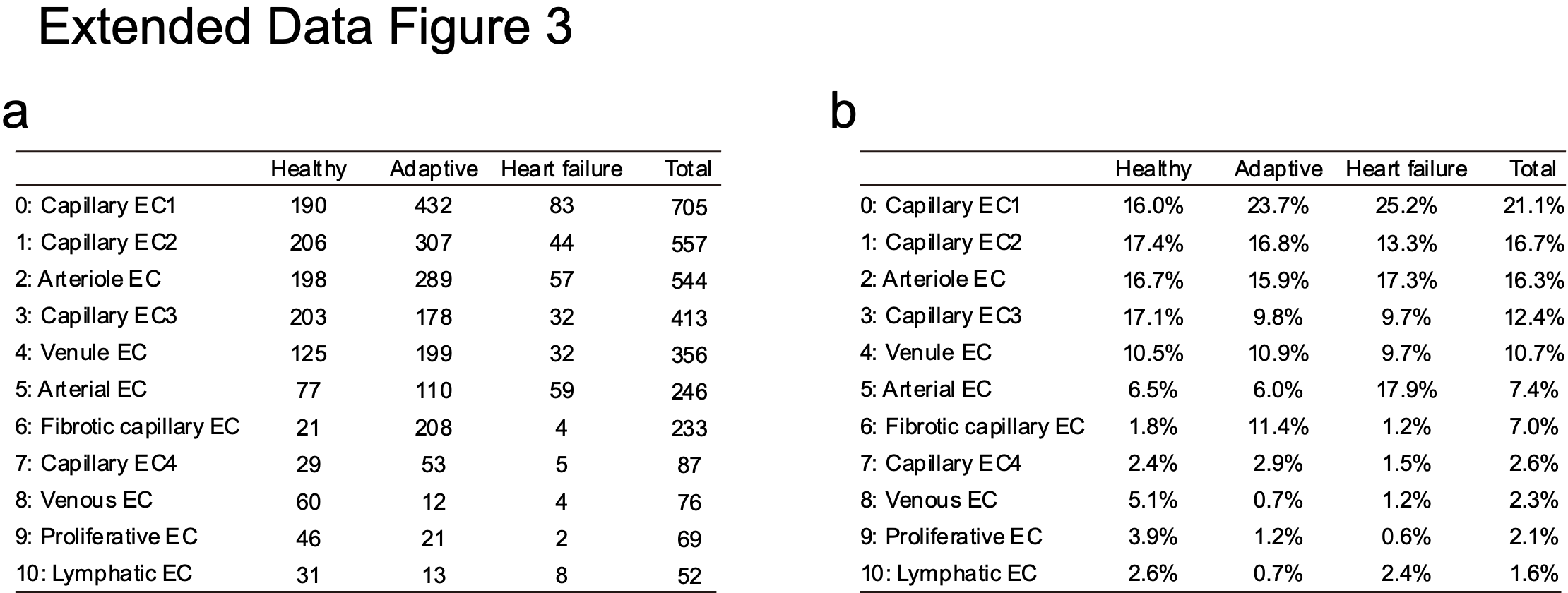

**Extended data fig.3: Detailed subtypes of endothelial cells.**

a-b, Annotated cell numbers used in these experiments. Rows were cell types. The columns represent the disease progression phases. Crude cell numbers (a) and proportions (b). EC, endothelial cells.

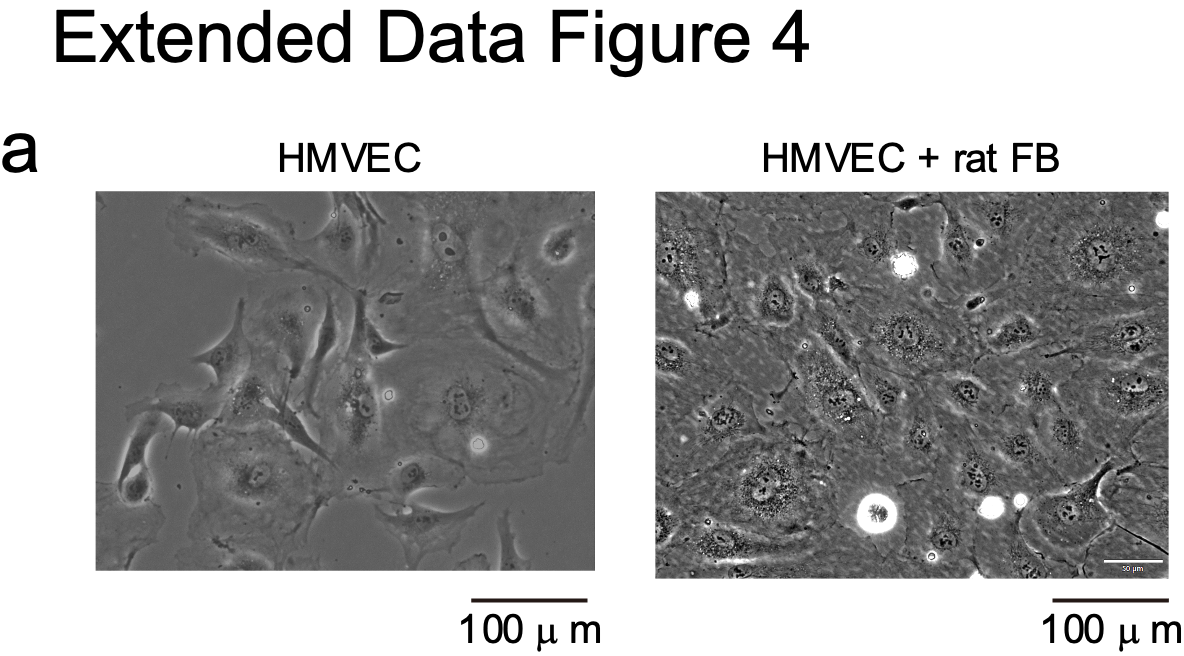

**Extended data fig. 4: Microscopic photos of co-cultured cells.**

a, Human microvascular endothelial cells (HMVEC, left) and co-cultured HMVEC/rat fibroblasts (right).

HMVEC, human cardiac microvascular endothelial cells; FB, fibroblasts.

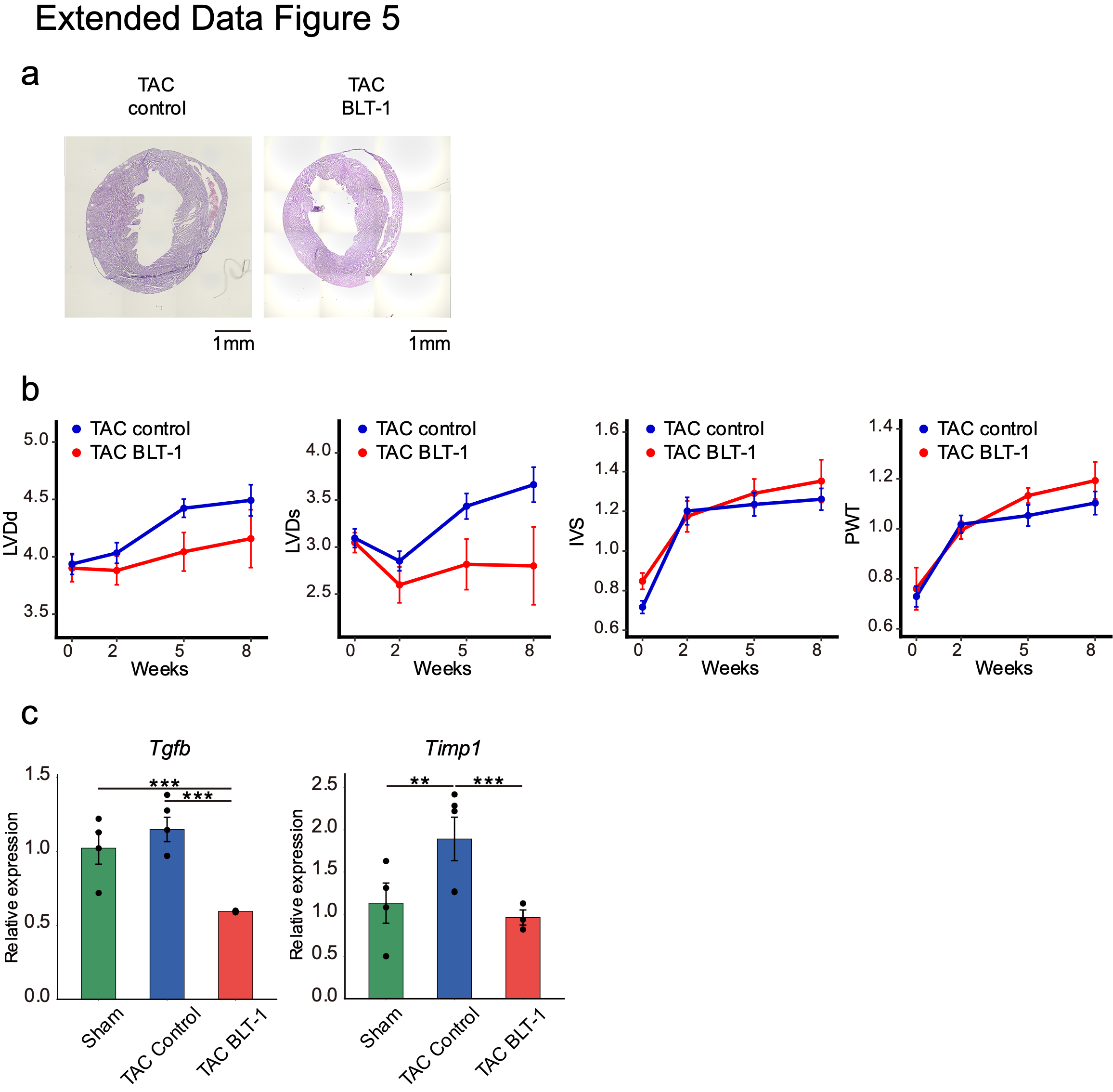

**Extended data fig. 5: Differences between control and BLT-1 treated TAC mice.**

a, Representative microscopic images of HE-stained control/BLT-1 treated TAC mice Control (left) and BLT-1 treated (right). b, Temporal trends in echocardiographic parameters in control/BLT-1 treated TAC mice. Blue lines represent control mice graphs and red lines represent BLT-1 treated mice. The x-axis represents weeks after TAC. The LVDd (leftmost), LVDs (second left), IVS (second right), and PWT (rightmost). The unit of the y-axis is millimeters. c, RNA expression profiles of Sham, TAC control and BLT-1 treated TAC mice (*n* = 9, 9, and 6, respectively). Y-axes are expressed relative to internal Gapdh. Green/blue/red bars represent Sham/TAC control/BLT-1 treated TAC mice. The dots represent raw data. Error bars represent the standard error. * indicates p < 0.05, ** indicates p < 0.01, and *** indicates p < 0.001. BLT-1, block lipid transport-1; TAC, transverse aortic constriction; HE, hematoxylin and eosin; LVDd, left ventricular end-diastolic diameter; LVDs, left ventricular end-systolic diameter; IVS, interventricular septum thickness; PWT, posterior wall thickness.

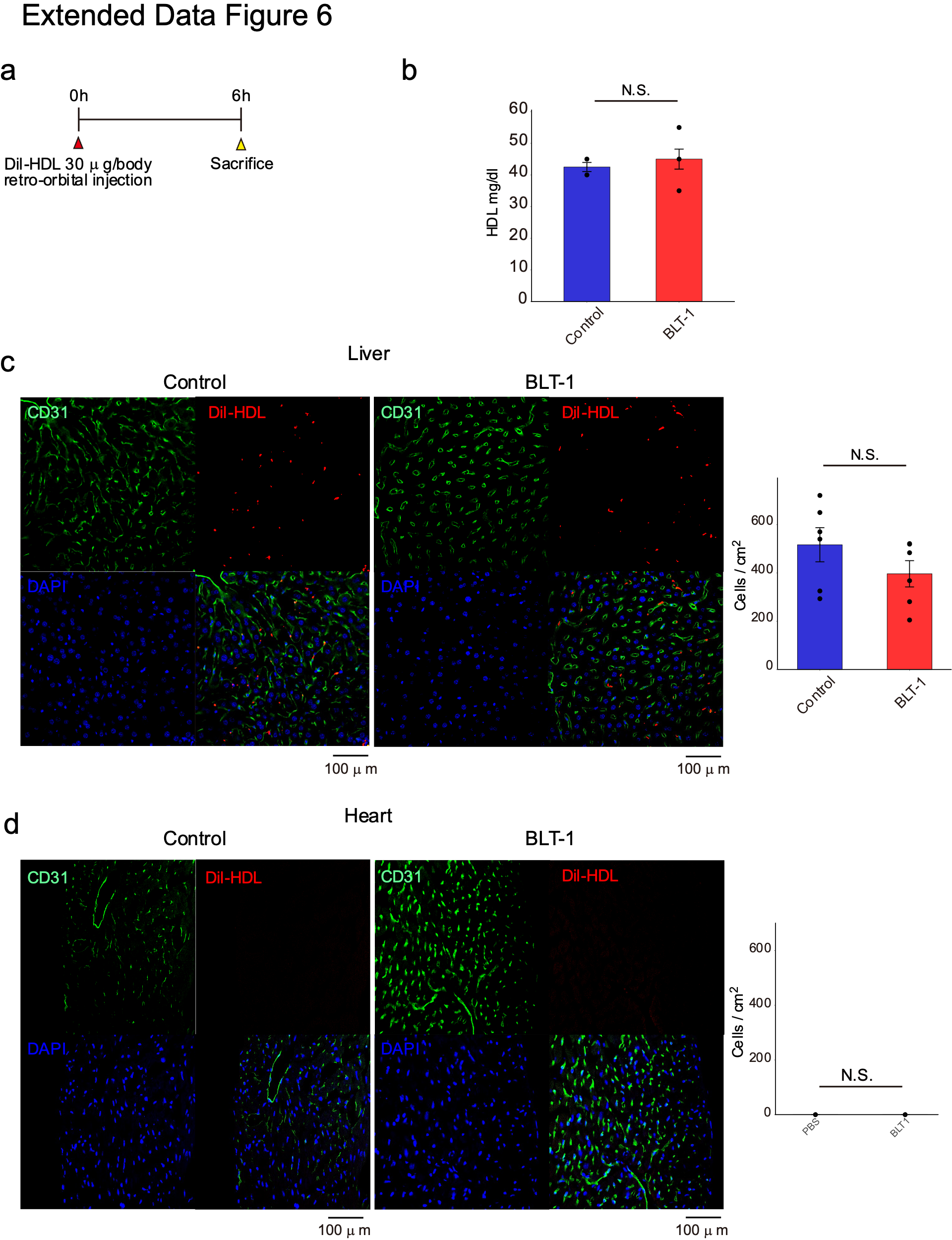

**Extended data fig. 6: HDL dynamics and BLT-1 effect in vivo.**

a, Experimental schematic of Dil-HDL injection to visualize in vivo HDL dynamics. Dil-HDL 30ug/body was retro orbitally injected to sample mice, and they were sacrificed after 6 hours. b, Serum HDL concentration in control and BLT-1 mice (*n* = 3). PBS 1 mL was intraperitoneally injected into the control mice for 7 days before sacrifice. BLT-1 1.25 mg DMSO/PBS was injected into the treatment group of mice in the same manner. c-d, Immunohistochemistry (IHC) images of control/BLT-1 treated mice and Dil-HDL-positive cell counts. Liver control IHC (c, left), BLT-1 treated IHC (c, middle), and Dil-HDL-positive cell counts (c, right). N.S. = “not statistically significant.” Control heart IHC (d, left), BLT-1 treated IHC (d, middle), and Dil-HDL-positive cell counts (d, right). No Dil-HDL-positive cells were detected in heart samples. HDL, high-density lipoprotein; BLT-1, block lipid transport-1; Dil-HDL, 1,1'- 3,3,3',3'-tetramethylindocarbocyanine perchlorate-labelled HDL; PBS, phosphate-buffered saline.

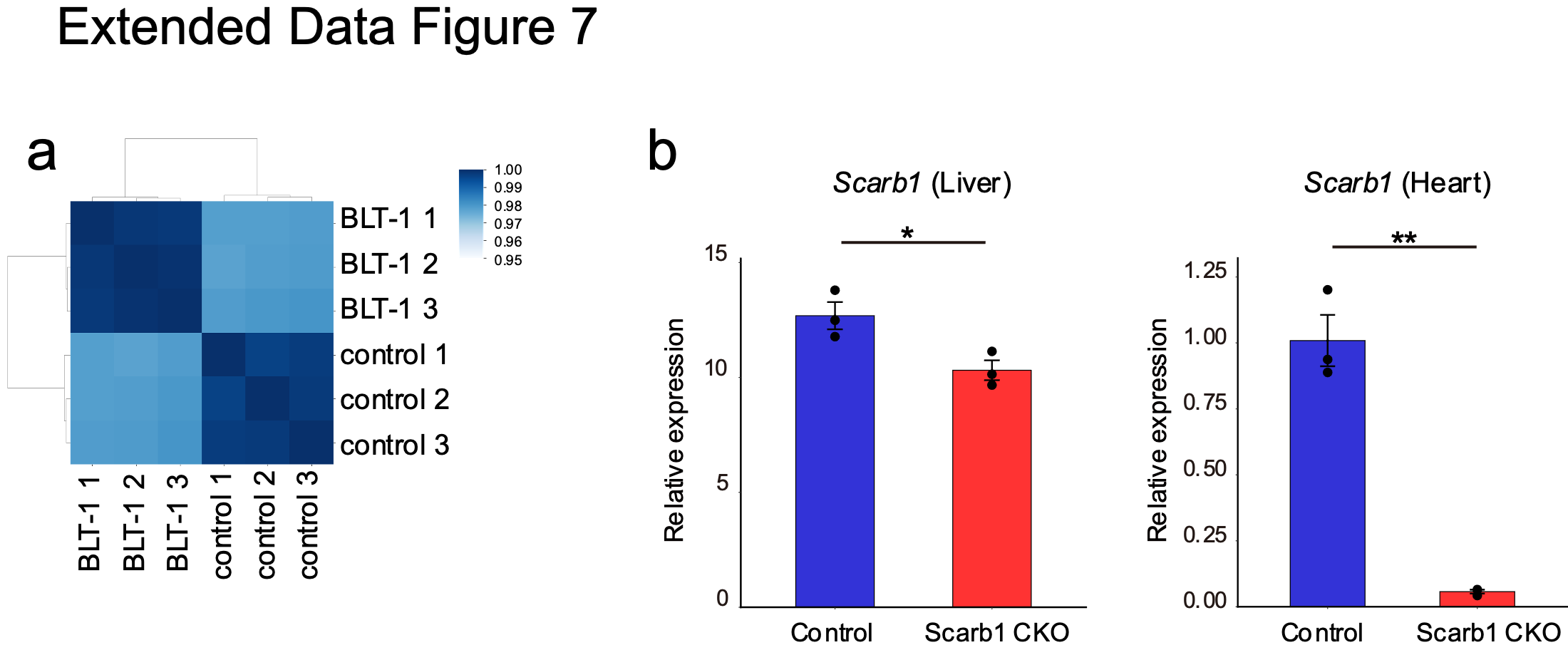

Extended data fig. 7: Effect of BLT-1 treatment on HMVEC and Scarb1 CKO confirmation.

a, Correlation matrix heatmap of control/BLT-1 treated HMVEC RNA expression. Darker blue indicates a stronger correlation. b, Scarb1 RNA expression profiles of control and Scarb1 CKO mice (*n* = 3). Y-axes are relative expression to internal Gapdh. Blue/red bars represent control/ Scarb1 CKO TAC mice. The dots represent raw data. Error bars represent the standard error. * indicates p < 0.05 and ** indicates p < 0.01. BLT-1, block lipid transport-1; HMVEC, human cardiac microvascular endothelial cells; RNA, ribonucleic acid; CKO, conditional knockout; TAC, transverse aortic constriction.

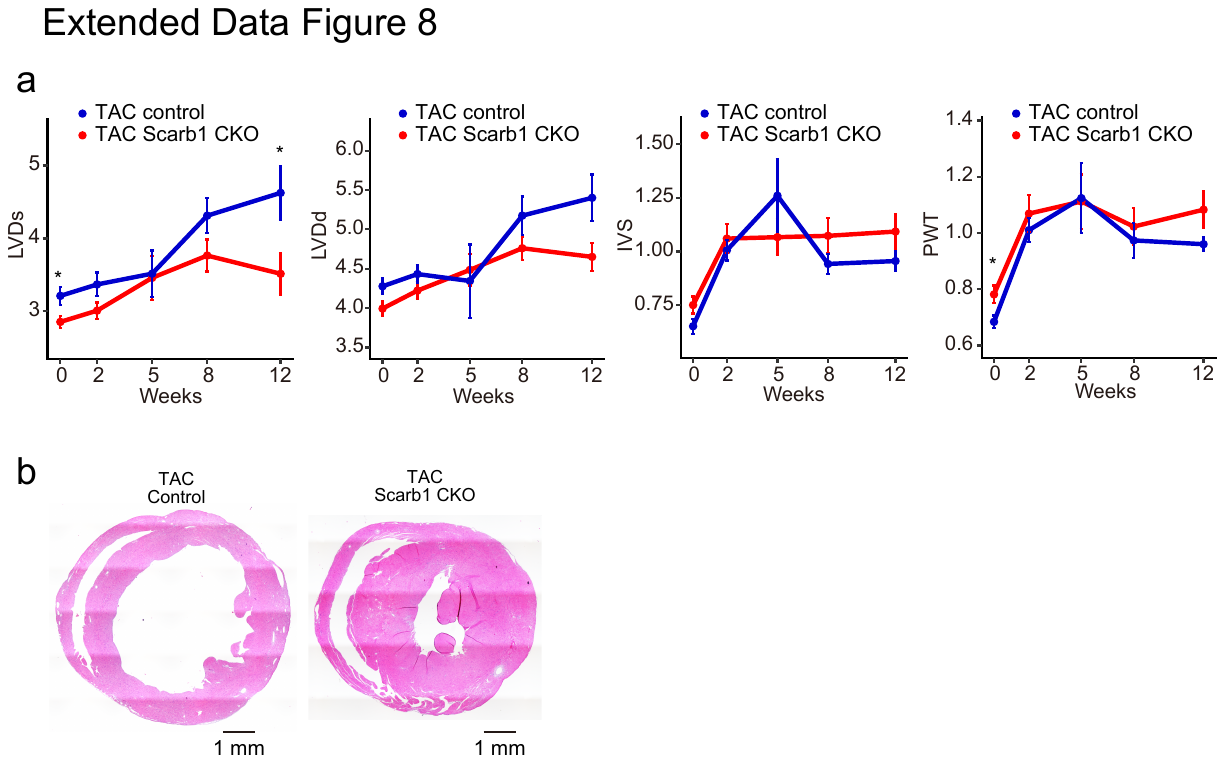

Extended data fig. 8: Differences between control and Scarb1 CKO TAC mice.

a, Temporal trends in echocardiographic parameters in control/Scarb1 CKO TAC mice. Blue lines represent control mice graphs, and red lines represent Scarb1 CKO. The x-axis represents weeks after TAC. LVDs (leftmost), LVDd (second left), IVS (second right), and PWT (rightmost). The unit of the y-axis is millimeters. * indicates p < 0.05. b, Representative microscopic images of HE-stained control/Scarb1 CKO TAC mouse specimens. Control (left) and Scarb1 CKO (right). CKO, conditional knockout; TAC, transverse aortic constriction.
